# Single cell fitness landscapes induced by genetic and pharmacologic perturbations in cancer

**DOI:** 10.1101/2020.05.08.081349

**Authors:** Sohrab Salehi, Farhia Kabeer, Nicholas Ceglia, Mirela Andronescu, Marc Williams, Kieran R. Campbell, Tehmina Masud, Beixi Wang, Justina Biele, Jazmine Brimhall, Jerome Ting, Allen W. Zhang, Ciara O’Flanagan, Fatemeh Dorri, Nicole Rusk, Hak Woo Lee, Teresa Ruiz de Algara, So Ra Lee, Brian Yu Chieh Cheng, Peter Eirew, Takako Kono, Jennifer Pham, Diljot Grewal, Daniel Lai, Richard Moore, Andrew J. Mungall, Marco A. Marra, IMAXT Consortium, Andrew McPherson, Alexandre Bouchard-Côté, Samuel Aparicio, Sohrab P. Shah

## Abstract

Tumour fitness landscapes underpin selection in cancer, impacting etiology, evolution and response to treatment. Progress in defining fitness landscapes has been impeded by a lack of timeseries perturbation experiments over realistic intervals at single cell resolution. We studied the nature of clonal dynamics induced by genetic and pharmacologic perturbation with a quantitative fitness model developed to ascribe quantitative selective coefficients to individual cancer clones, enable prediction of clone-specific growth potential, and forecast competitive clonal dynamics over time. We applied the model to serial single cell genome (*>*60,000 cells) and transcriptome (*>*58,000 cells) experiments ranging from 10 months to 2.5 years in duration. We found that genetic perturbation of *TP53* in epithelial cell lines induces multiple forms of copy number alteration that confer increased fitness to clonal populations with measurable consequences on gene expression. In patient derived xenografts, predicted selective coefficients accurately forecasted clonal competition dynamics, that were validated with timeseries sampling of experimentally engineered mixtures of low and high fitness clones. In cisplatin-treated patient derived xenografts, the fitness landscape was inverted in a time-dependent manner, whereby a drug resistant clone emerged from a phylogenetic lineage of low fitness clones, and high fitness clones were eradicated. Moreover, clonal selection mediated reversible drug response early in the selection process, whereas late dynamics in genomically fixed clones were associated with transcriptional plasticity on a fixed clonal genotype. Together, our findings outline causal mechanisms with implication for interpreting how mutations and multi-faceted drug resistance mechanisms shape the etiology and cellular fitness of human cancers.

## Introduction

Cellular fitness underpins the tissue population dynamics of cancer progression and treatment response. Yet, quantifying fitness in heterogeneous cell populations, and identifying causal mechanisms shaping fitness landscapes remain as open problems, impeding progress in developing effective and durable therapeutic strategies. In particular, quantitative fitness modeling of cancer cells has numerous and diverse implications; attributing clonal dynamics to drift or selection, identifying the determinants of clonal expansion, enabling causal inference, and forecasting growth trajectories. Drug resistance and etiology are among the key unresolved areas of investigation that require advanced understanding of fitness in cancer. For example, drug resistance mechanisms are commonly attributed to phenotypic plasticity encoded via epigenetic changes^1, 2^, or evolutionary selection of pre-existing genomic clones^3^. However, the relative contribution of these processes when studied in tandem is poorly understood and requires integrated genome-transcriptome investigation. Moreover, how changes in genomic architecture brought about by copy number alterations drive etiologic processes remains understudied. Perturbations to induce fitness changes include genetic editing with cancer gene mutations, and pharmacologic drug selection to profile kinetics of therapeutic response. We contend that quantitatively ascribing fitness values to clonal dynamics over long-range timeseries in the context of such perturbations would provide higher order insights in etiology and drug resistance than current models allow.

Previous work has established models of fitness through interpreting allelic measurements of single snapshots^4–8^ from bulk sequencing over large patient cohorts^9^, timeseries study of cell free DNA^10^, multiregion sequencing^5, 11–14^ and estimating fitness landscapes of clonal haematopoiesis^15^. However, the cancer field has generally lacked serial measurements from patient derived tissues to directly observe cancer evolution over realistic timescales. This has impeded a thorough understanding of causal factors driving selection, achieved in other biological systems through studying granular timeseries with population genetics modeling^16^. The majority of work in cancer has focused on bulk tumour sequencing, where cellular population structure decomposition approaches are limited. Single cell genome measurements to scalably define clonal populations in cancer over thousands of cells have only recently emerged^17, 18^, enabling identification of rare populations, precise tracking of clones and robust clone-specific measurements suitable for population genetics modeling. Combined with single cell transcriptome measurements, dissecting concomitant genotype-phenotype contributions to clonal dynamics at granular timescales is now tractable.

Here we set out to establish quantitative fitness attributes of cancer cells as predictive measures of their growth potential in polyclonal systems. We used scaled single cell genome sequencing (*>*60,000 cells) using the direct library preparation (DLP+) method ^19^, predicted clonal population trajectories, and defined clonal fitness landscapes. We combined these results with scRNAseq on the same samples (*>*58,000 cells) to determine concomitant and divergent changes at the phenotypic level. Studying timeseries data of serially passaged mammary epithelial cells and patient derived breast cancer xenografts over multi-year timescales, our results reveal how genomic copy number architectures shape fitness landscapes, how predicted selective coefficients can forecast clonal competition dynamics and how pharmacologic perturbation inverts pre-treatment fitness landscapes. Our work has implications in at least three areas: 1) predicting evolution in cancer; 2) understanding how genomic instability processes confer fitness; 3) parsing the long term kinetics of drug resistance into phases of clonal evolution and transcriptional plasticity.

## Results

### Modeling clonal fitness and selection

We developed an experimental and computational platform consisting of three major components: scaleable phylogenetics for single cell genomes to identify clones, timeseries sampling of immortal cell lines (*>*10 months) and patient derived xenografts (*>*2.5 years) to observe clonal dynamics, and a mathematical model for inferring clone-specific fitness measures (**Fig. 1**). For normal human breast epithelial cells^20^ *in vitro* and in breast cancer PDX^21, 22^ (**Fig. 1**A), we sequenced *>*60,000 cells over interval passaging (**Fig. 1B**, **Supplementary Table 1**) with single cell whole genome sequencing^17^ (scDNAseq) measuring single cell copy number profiles and computing phylogenetic trees to identify genotypic clones and their relative abundances as a function of time. Genetic (p53 biallelic inactivation for cell lines) and pharmacologic (cisplatin dosing in PDX models) perturbations were applied to determine their impact on fitness landscapes. In conjunction with scDNAseq, we generated scRNAseq from *>*58,000 cells to establish clone-specific phenotypes (**Fig. 1C**, **Supplementary Table 2**), estimated through clone-specific expression profiles. Timeseries clonal abundance observations were modeled using an implementation of the Wright-Fisher diffusion process (**Fig. 1D**) we called `fitClone` (Supplementary Information). `fitClone` simultaneously estimates growth trajectories, *X*_*i*_ and fitness coefficients, *s*_*i*_ for each clone *i* in the population (**Fig. 1E**). We note that increasing 1 + *s*_*i*_ indicates positive selection and higher growth potential. The model accounts for drift as well as selection, with fitness estimated relative to a reference population, where *s* = 0 by construction. As a generative process, the model can be used for forecasting evolutionary trajectories of specific clones.

**Figure 1.**
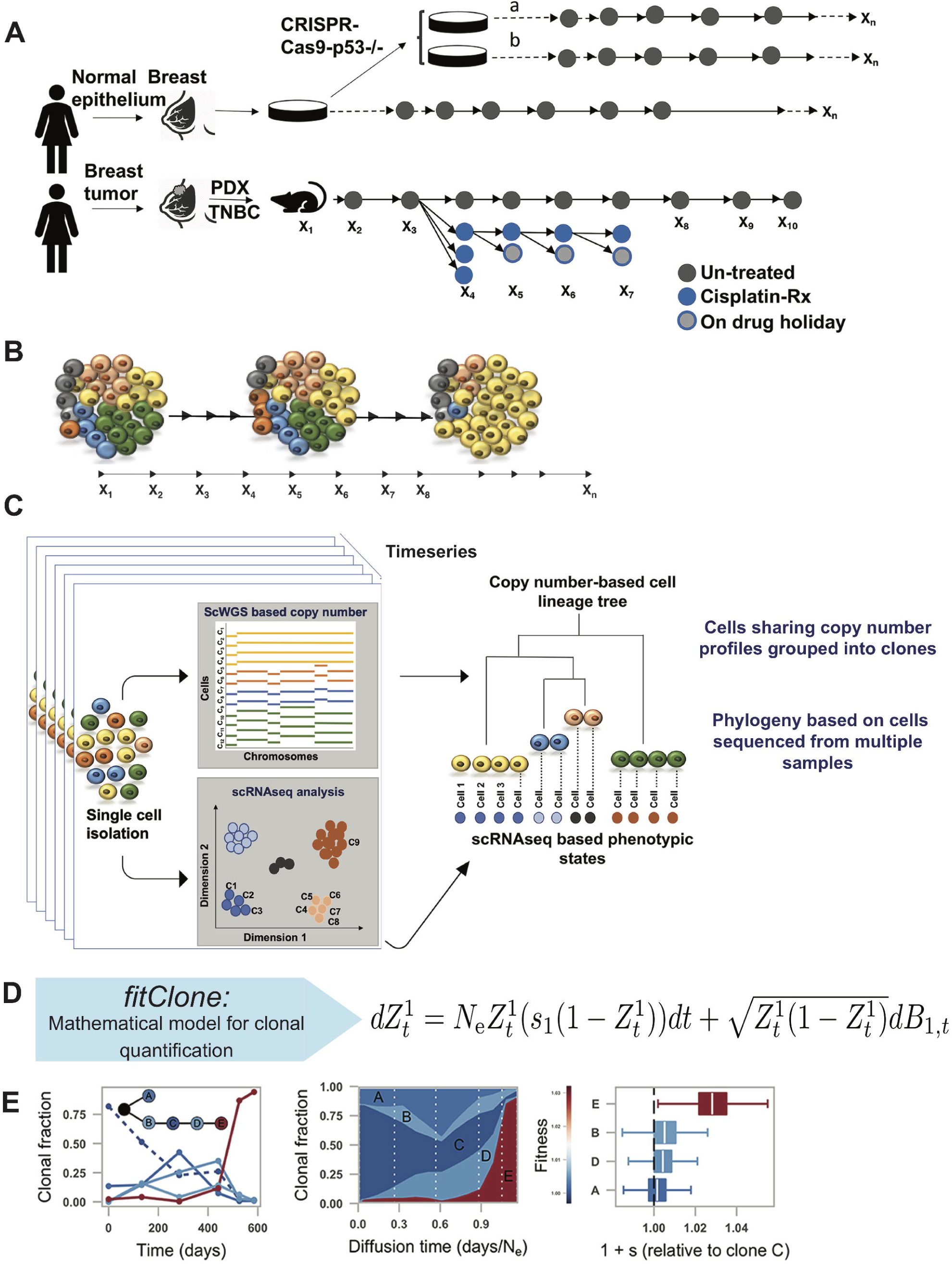
Schematic overview of experimental design for quantitatively modeling clone-specific fitness. **A)** Timeseries sampling from in vitro or PDX systems **B)** Clonal dynamics of cell populations observed over time **C)** Whole genome and transcriptome single cell sequencing of timeseries samples and shared phylogenetic tree inference **D)** `fitClone`: mathematical modeling of fitness with diffusion approximations to the K-type Wright-Fisher model **E)** Application of `fitClone` to previously published clonal dynamics in PDX showing observed data (left), inferred trajectories (middle) and posterior distributions of fitness coefficients (right).

We carried out simulation experiments to establish the theoretical behaviour and limitations of `fitClone`, over a range of parameters. Model fits were robust to effective population size and number of clones (**Supplementary Fig. 1B,C**). In two key advances, simultaneous modeling of multiple clones was superior to modeling each clone independently, and accounting for stochasticity in the dynamics via the diffusion process led to more accurate selection coefficient estimates than with a deterministic growth model (**Supplementary Fig. 1A**). Together these simulations established a rationale for systematic modeling of all clones in a unified approach with a generative process. We then applied `fitClone` to previously published data, wherein experimentally derived reproducible clonal dynamics had been reported in breast cancer PDXs^21^. From one of these lines, the abundance of five major clones (A-E, **Supplementary Table 3**), as determined by single cell genotyping was measured over six serial passages. One clone (clone E) was found in the original study to undergo a selective sweep in repeat and independent passaging^21^, suggesting selection due to higher fitness. `fitClone` estimates converged with Clone E bearing the highest fitness (1 + *s*=1.03 ± 0.01), consistent with positive selection over the timeseries (**Fig. 1E**), and thus representing a proof of principle application of `fitClone` on real-world data.

### Segmental aneuploidies drive positive selection in diploid p53 deficient cells

We next applied our framework to immortalized 184hTert diploid breast epithelial cell lines^20^ to determine mechanisms by which *TP53* mutation induces clonal expansions and fitness trajectories. *TP53* is the most abundantly mutated gene in all human cancers^23^, and specifically in breast cancers^7, 24^. Known to be permissive of genomic instability, *TP53* loss is often acquired early in evolution and results in profound alteration of the copy number landscape^14, 25–27^. We therefore asked whether specific clonal expansions could be observed, and moreover if selective fitness advantages could be quantified, as a function of *TP53* ablation, thereby modeling copy number driven etiologic processes in a controlled system of immortalized mammary epithelial cells. *TP53* wildtype (*p53 WT*) timeseries sampling (60 passages over 300 days, 4 samples) was contrasted with two isogenic *TP53* deficient `NM_000546(TP53):c.[156delA];[156delA]`^17^ parallel branches (*p53-/-a* and *p53-/-b*), each passaged over 60 generations (285 and 220 days, respectively) and sampled 7 times. A median of 1,231 cells per passage were sequenced yielding a total of 6620, 7935, 9615 single cell genomes for each timeseries, respectively (**Supplementary Fig. 2A,B**, **Supplementary Table 1**). For each of *p53 WT, p53-/-a, p53-/-b* we inferred single cell copy number profiles, constructed a phylogenetic tree to establish clonal lineages (Supplementary Information, **Supplementary Fig. 3**) and measured clonal abundances as a function of time (**Fig. 2A-C**). Modeling the abundances with `fitClone` (**Supplementary Tables 4** and **5**) revealed *p53 WT* clonal trajectories consistent with small differences over the posterior distributions of fitness coefficients amongst four major clones (**Fig. 2D**). In contrast, p53 mutant branches each showed significant expansions of clones with aneuploid genotypes, where the founder diploid population was out-competed. Relative to *p53 WT*, rates of expansion of p53 mutant, aneuploid clones were significantly higher, leading to rapid depletion of diploid cells (**Fig. 2D**, p=6.72e-04). Pairwise difference of selective coefficients *s*_*i*_ − *s*_*j*_ (Δs) between clones in the `fitClone` inference process was larger in the *p53-/-* lines relative to *p53 WT* (**Fig. 2E**). This suggests that p53 mutation permits expansion of clones at higher rates, and that these clones have measurably higher positive selection coefficients.

**Figure 2.**
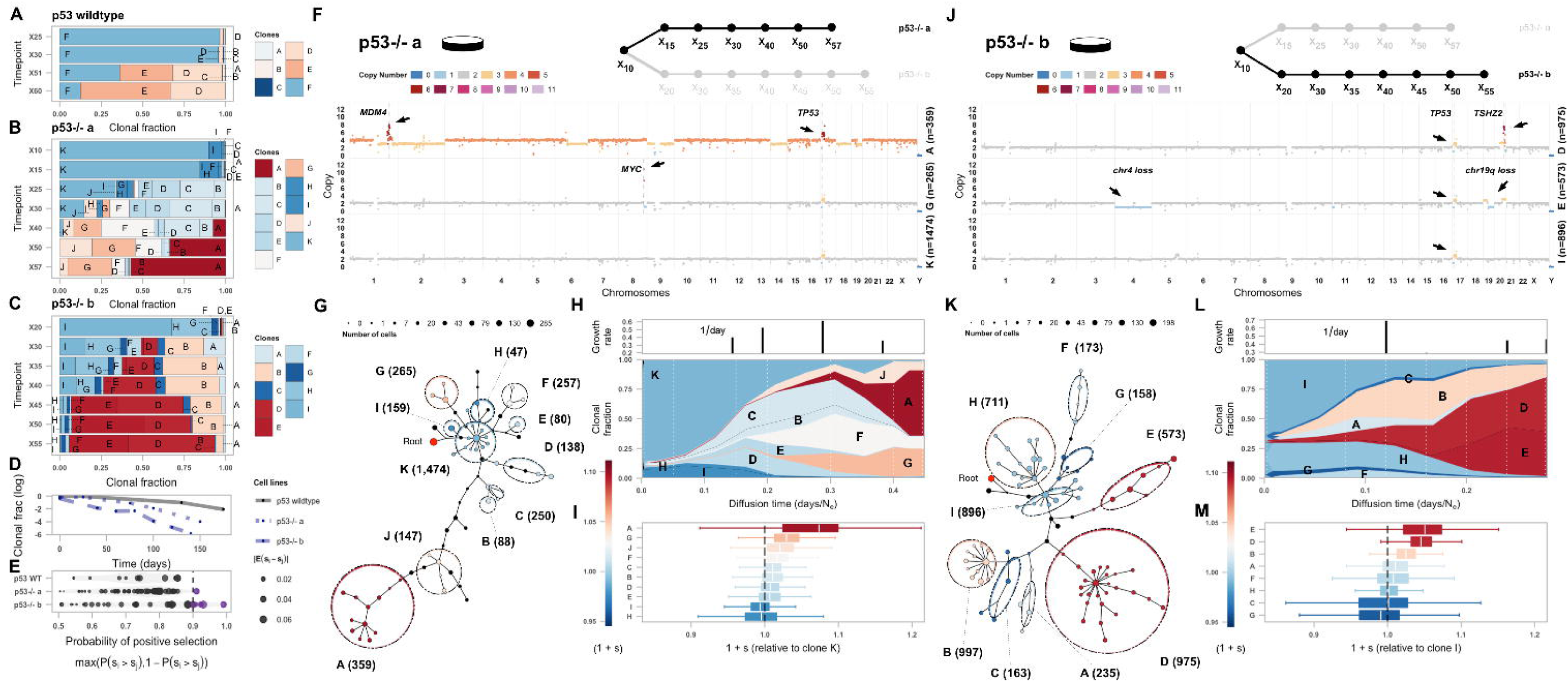
Impact of p53 mutation on modes of selection on 184hTert cells. **A-C)** Clonal dynamics of p53 wildtype, and two independent timeseries of p53 mutant 184hTert mammary epithelial cell lines **D)** Clonal fraction of diploid reference over time **E)** Distribution over magnitude of difference between selective coefficients of pairs of clones **F)** Clonal genotypes of 3 representative clones for *p53-/-a* showing high level amplification of *MDM4* in clone A and *MYC* in clone G. Reference diploid clone K shown for comparison **G)** Phylogeny of cells over the timeseries *p53-/-a* where nodes are groups of cells (scaled in size by number) with shared copy number genotype and edges represent distinct genomic breakpoints **H)** Inferred trajectories and **I)** selective coefficients of `fitClone` model fits to *p53-/-a* **J)** Clonal genotypes of 3 representative clones for *p53-/-b* showing high level amplification of *TSHZ2* in clone D, Chr4 loss in clone E. Reference diploid clone I shown for comparison. **K-M** Analogous to **F-I** but for *p53-/-b*.

**Figure 3.**
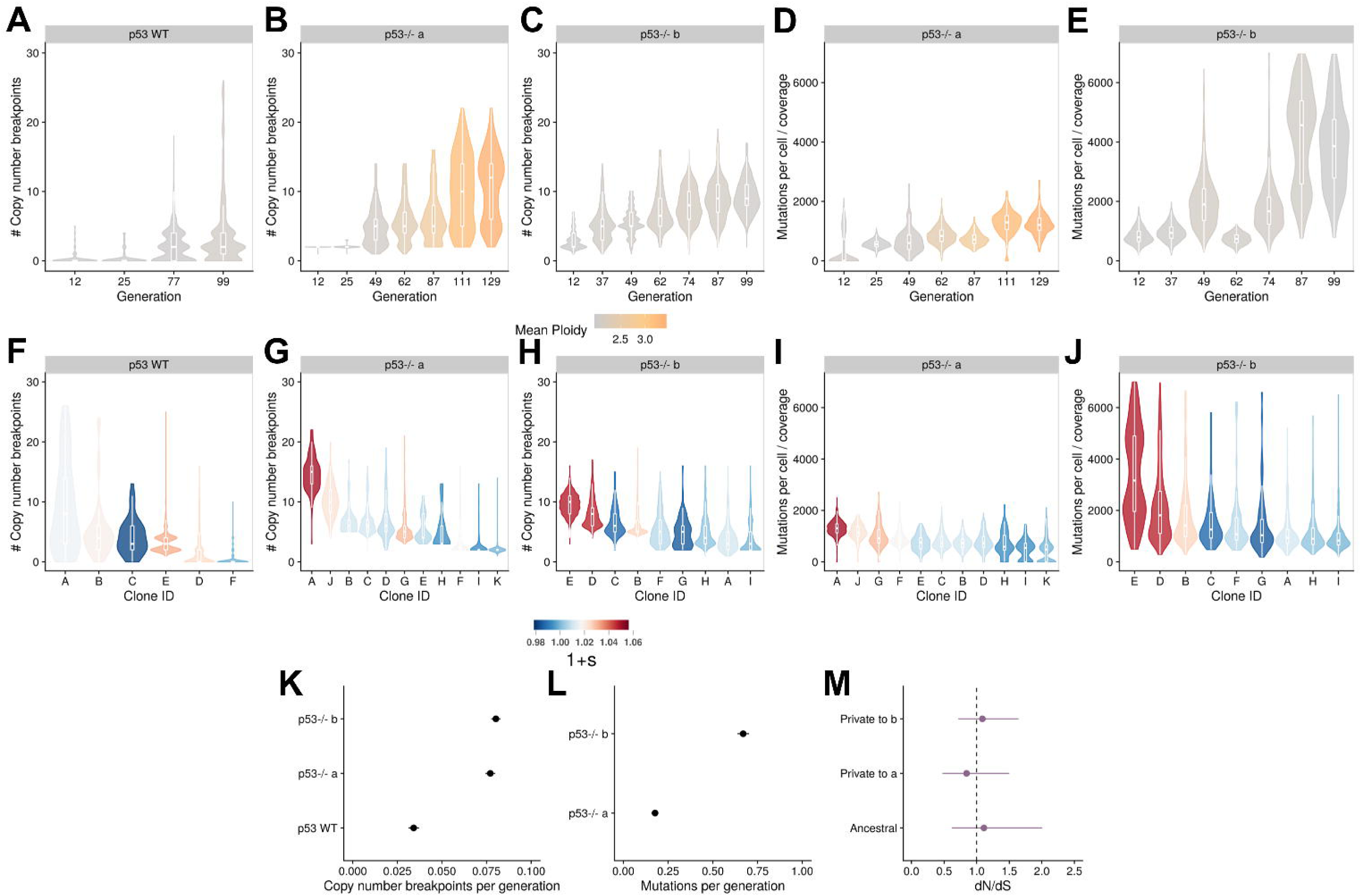
Structural variant and mutation rates of 184hTert cells. Distribution over copy number breakpoints/cell as a function of generation for **A)** *p53 WT* **B)** *p53-/-a* **C)** *p53-/-b*. Distribution over point mutations/cell as a function of generation for **D)** *p53-/-a* and **E)** *p53-/-b*. Clone specific distributions over copy number breakpoints/cell, colored by fitness coefficients for **F)** *p53 WT* **G)** *p53-/-a* and **H)** *p53-/-b*. Clone specific distributions over point mutations/cell, colored by fitness coefficients for **I)** *p53-/-a* and **J)** *p53-/-b*. Copy number breakpoints **K)** and mutations **L)** per generation for *p53 WT, p53-/-a* and *p53-/-b*. dN/dS ratios for mutations shared between *p53-/-a* and *p53-/-b* and unique to both lines.

The p53 mutant lines harbored 11 (size range 47 to 1,474 cells, median 204), and 10 (size range 158 to 997 cells, median 404) distinct clones for *p53-/-a* and *p53-/-b*, respectively (**Supplementary Table 3**). In each series the diploid founder clones, devoid of detectable copy number alterations, were systematically out-competed by populations that had acquired at least one copy number alteration (**Fig. 2F-M**). Some similarities in copy number events were observed during the two replicate timeseries. These included gains in chromosomes 13, 19p and 20, and losses on chromosomes 8p and 19q (**Supplementary Fig. 3**). However, by the end of the timeseries, the genotypes in the two lines had diverged considerably. Selective coefficients were highest in clones with focal amplifications of known prototypic oncogenes in breast cancer^7, 24, 27, 28^ including in *MDM4, MYC* (**Fig. 2F**) and *TSHZ2* and (**Fig. 2J**), in some cases on a whole genome doubled background. Clone A, the highest fitness clone in *p53-/-a* (57% of cells at last timepoint, 1 + *s*=1.05 ± 0.09) exhibited a whole genome doubling event (18 chromosomes with four copies) and harboured a focal, high level amplification at the *MDM4* locus on Chr1q (**Fig. 2F**). Clone G (27% of cells at last timepoint, 1 + *s*=1.03 ± 0.03), the next highest fitness clone in *p53-/-a* remained diploid, with the exception of a focal high level amplification precisely at the *MYC* locus on Chr8q (**Fig. 2G**). By contrast Clone K, here chosen as the reference clone for modeling, remained entirely diploid and exhibited a monotonically decreasing trajectory (from 90% to 0% of cells over the timeseries, **Fig. 2H,I**). In *p53-/-b*, two clones exhibited non-neutral, positive selective coefficients (**Fig. 2J**). Clone D (52% of cells at last timepoint, 1+*s*=1.05 ± 0.02) harbored a Chr20q single copy gain with an additional high level amplification at the *TSHZ2* locus, while Clone E (35% of cells at last timepoint, 1 + *s*=1.05 ± 0.04) harbored a Chr4 loss, Chr19p gain/19q loss and Chr20q single copy gain (**Fig. 2F**). As seen in *p53-/-a*, the ‘root’ Clone I that remained diploid was systematically outcompeted, diminishing from 68% to 0% abundance over the timeseries (**Fig. 2K,L**).

We next tested whether the genomes of individual cells became progressively more aberrant over time and whether this correlated with measurements of clonal fitness. We estimated both sample and clone specific mutation rates at the copy number breakpoint and point mutation level (as previously described^17^). Both *p53-/-a* and *p53-/-b* exhibited increased mutations and breakpoints over time relative to *p53 WT* (**Fig. 3A-E**). Cells accumulated 0.08 additional breakpoints (p*<*0.0001) and an average of 0.4 additional mutations (0.17 in *p53-/-a* and 0.67 in *p53-/-b*, p*<*0.001) per generation (**Fig. 3K,L**, linear regression model, Supplementary Information), while the *p53 WT* line accumulated 0.03 additional breakpoints per generation (**Fig. 3K**). As this cell line was used as the reference for clone-specific point mutation detection in *p53-/-* cell lines, by definition no *p53 WT* point mutations were analysed. Clone level distributions of breakpoints and mutations were positively correlated with inferred fitness coefficients in both *p53-/-* lines (**Fig. 3G-J**, p=0.001) but not in *p53 WT* (**Fig. 3F** linear regression, p=0.5). Point mutation analysis in individual cells did not reveal any putative driver mutations associated with the fittest clones, selection pressure of mutations as measured by the dN/dS ratio^9^ were not increased (**Fig. 3M**), suggesting SNVs were not driving clonal expansions in this system.

These results indicate that the impact of genetic perturbation on selection can be measured and modeled in clonal populations. In particular, p53 mutation, known to be an early event in the evolution of many cancers^29^, yields clonal expansions driven by whole genome, chromosomal and segmental aneuploidies, conferring quantitative fitness advantages over cells that maintain diploid genomes. Orthogonal analysis on mutation rates was consistent with expected increases as a function of inferred fitness coefficients, further corroborating biological relevance of `fitClone` estimates. Modes of positive selection involving high level amplification of proto-oncogenes often seen in human breast cancer, and aneuploidies in general, suggest that in vitro genetic manipulations can induce fitness-enhancing genomic copy number changes consistent with etiologic roles in cancer.

### Clone-specific fitness estimates forecast clonal competition trajectories

We next modeled clonal expansions observed during serial passaging of *TP53* mutant PDX tumours to determine the predictive capacity of fitness coefficients (**Fig. 4, Supplementary Table 3**). We generated single cell genomes from 8 serial PDX transplants over 721 days from a HER2 positive (HER2+) breast cancer with a *TP53* p.A159P missense mutation, and contrasted this with 10 serial samples over 1002 days from a triple negative breast cancer (TNBC) PDX with a *TP53* p.R213* non-sense mutation (**Supplementary Fig. 2C,E**, **Supplementary Tables 6** and **7**). A median of 907 single cell genomes were sequenced per passage for a total of 11,705 and 10,553 single cell genomes from the HER2+ and TNBC series, respectively (**Supplementary Table 1**). Both series exhibited progressively higher tumour growth rates over time (**Fig. 4F,J**, **Supplementary Fig. 2D**). Data were analysed as per the *in vitro* lines described above and modeled with `fitClone` (**Supplementary Tables 4** and **5**). The HER2+ series exhibited 4 distinct clones ranging in size from 134 to 1,421 cells (median 319, **Fig. 4E**), and the TNBC series exhibited 8 distinct clones with 18 to 680 cells (median 556, **Fig. 4I**). In the HER2+ model, clonal trajectories were consistent with selective coefficients with small relative differences in fitness (**Fig. 4F,G**, mean = 1.01 ±0.01, Supplementary methods). By contrast, the TNBC model trajectories resulted in a high positive selective coefficient for a minority of clones (**Fig. 4J,K**, mean = 1.03 ±0.11, Supplementary Information). Consistent with increased dynamics in the TNBC series, we found an initial increase of 0.1 breakpoints per cell per generation in the first 4 passages (linear regression, p=0.03, *R*^2^=0.49, **Supplementary Fig. 4A**). After this initial increase the average number of breakpoints per cell remained constant (linear regression, p=0.8, *R*^2^=0.08). In the HER2+ line we observed a small decrease of 0.04 copy number breakpoints per generation (linear regression, p=0.002, *R*^2^=0.05). We note that Clone E in TNBC swept the population over the last 3 timepoints (**Fig. 4J,K,** n=541 over the timeseries). Clone E had the highest selective coefficient (1 + *s*= 1.08 ± 0.043), having grown from undetectable proportions in earlier timepoints to 58% of cells by the end of the timeseries. Clone E also had the highest number of breakpoints with 12.8 additional copy number breakpoints per cell, relative to the reference clone C with the lowest (linear regression with coverage breadth, ploidy and cell cycle state as covariates, p *<*0.0001, *R*^2^=0.364, **Supplementary Fig. 4C**).

**Figure 4.**
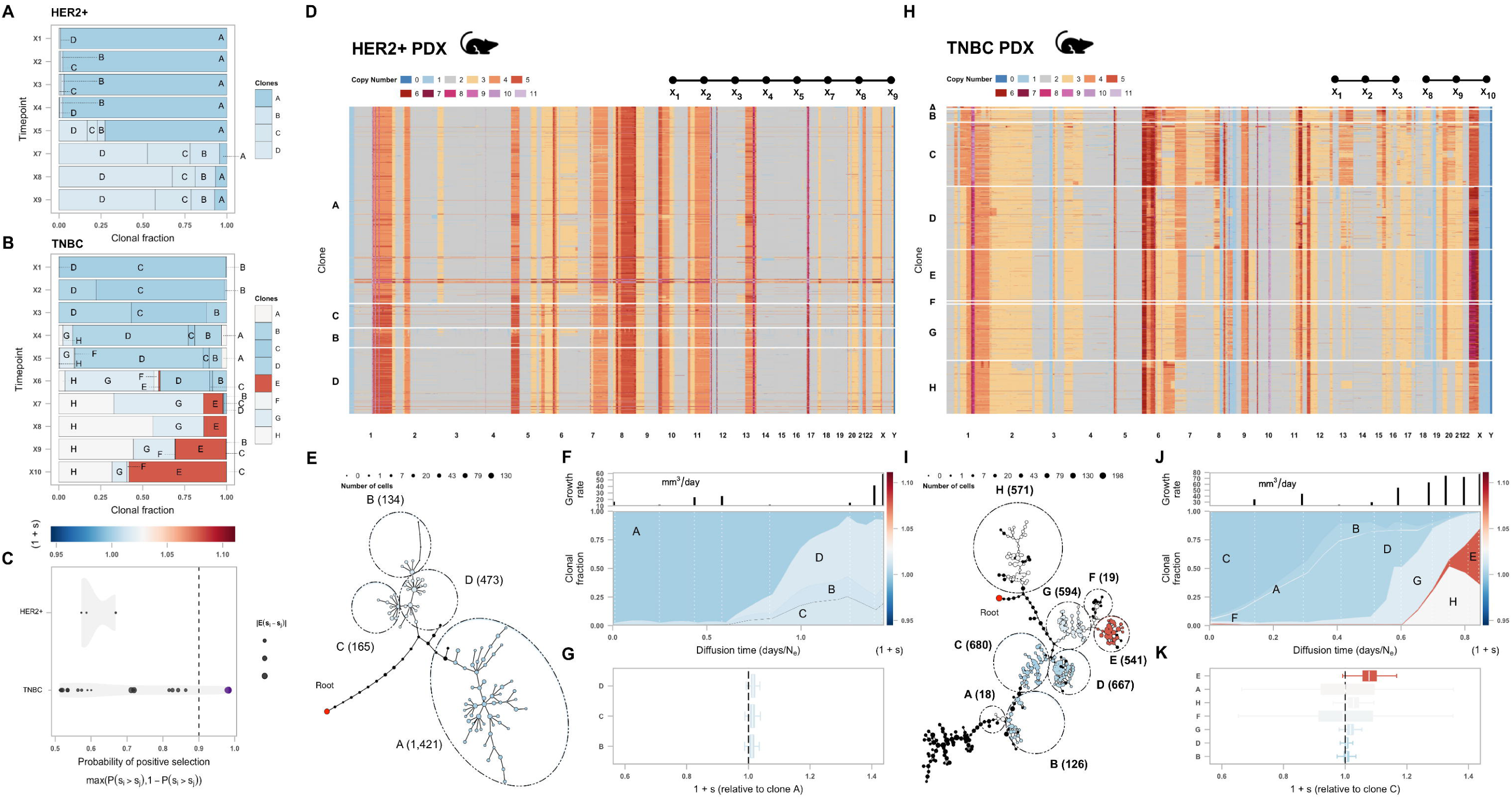
Comparison of fitness landscapes of breast cancer PDX models. **A-C)** Clonal dynamics of HER2+ and TNBC models (clones colored by inferred fitness coefficients). **D)** Heatmap representation of copy number profiles of 2,193 cells, grouped in 4 phylogenetic clades. **E)** Phylogeny as per **Fig. 2** for HER2+. **F)** Inferred `fitClone` trajectories and **G)** selective coefficients for the HER2+ model. **H-K)** Analogous plots for the TNBC model (n=3,216 cells).

We next asked whether the high predicted fitness of Clone E was a true indicator of positive selection through a physical clonal mixing and re-transplant experiment. Enforced clonal competition of higher fitness clones with lower fitness counterparts should result in re-emergence or fixation of high fitness clones, even when re-starting from a low population prevalence. To test this, we forward-simulated trajectories from `fitClone` using the estimated selective coefficients (A=1.03 ± 0.16, B=1.00 ± 0.01, C=0.00, D=1.00 ± 0.01, E=1.10 ± 0.10, F=1.01 ± 0.13, G=1.01 ± 0.02, H=1.02 ± 0.03) and starting clonal proportions of (A=0.00, B=0.07, C=0.25, D=0.51, E=0.02, F=0.00, G=0.08 H=0.07), derived by physically mixing cells from a late (X8) and an early (X3) passage of the TNBC series (**Supplementary Fig. 5A**, **Fig. 5A,B**). We note that `fitClone` assumes non-zero initial values for all clones, permitting growth of even exceedingly rare clones (Supplementary Information). We generated 47,000 trajectories from the model (**Fig. 5A**) and found Clone E with the highest probability of fixation (0.32), followed by Clones A (0.17) and F (0.12). Fixation probabilities for the remainder of clones were low (*<*0.01 for H and D) or zero (for B, C, and G). We experimentally tested these predictions by initiating a new PDX with the remixed population, serially passaged over 4 timepoints (**Fig. 5B**, **Supplementary Fig. 5A**), and sequenced with DLP+ (7,839 single cell genomes, median 1,354.5 per library). Seven clones from the original timeseries were recapitulated in the mixture timeseries (all but Clone A) with between 26 to 499 (median 162) cells (**Fig. 5C,D**). Clones with higher selective coefficients swept through the mixture timeseries by passage 4 (**Fig. 5D,E**). Comparison of model fits of the original and mixture timeseries yielded similar posterior distributions for the majority of clones (**Fig. 5F,G**). While Clone A was not detected in the mixture, Clone F emerged as a high fitness clone (1+*s*=1.10 ± 0.07). Notably, F was phylogenetically proximal to Clone E (**Fig. 5C**) and thus likely represented a biologically similar population. In the last timepoint, the clade composed of clones E and F comprised 94% of cells, outcompeting low-fitness clones and thus recapitulating `fitClone` coefficients predictions and forecasts of clonal competition.

**Figure 5.**
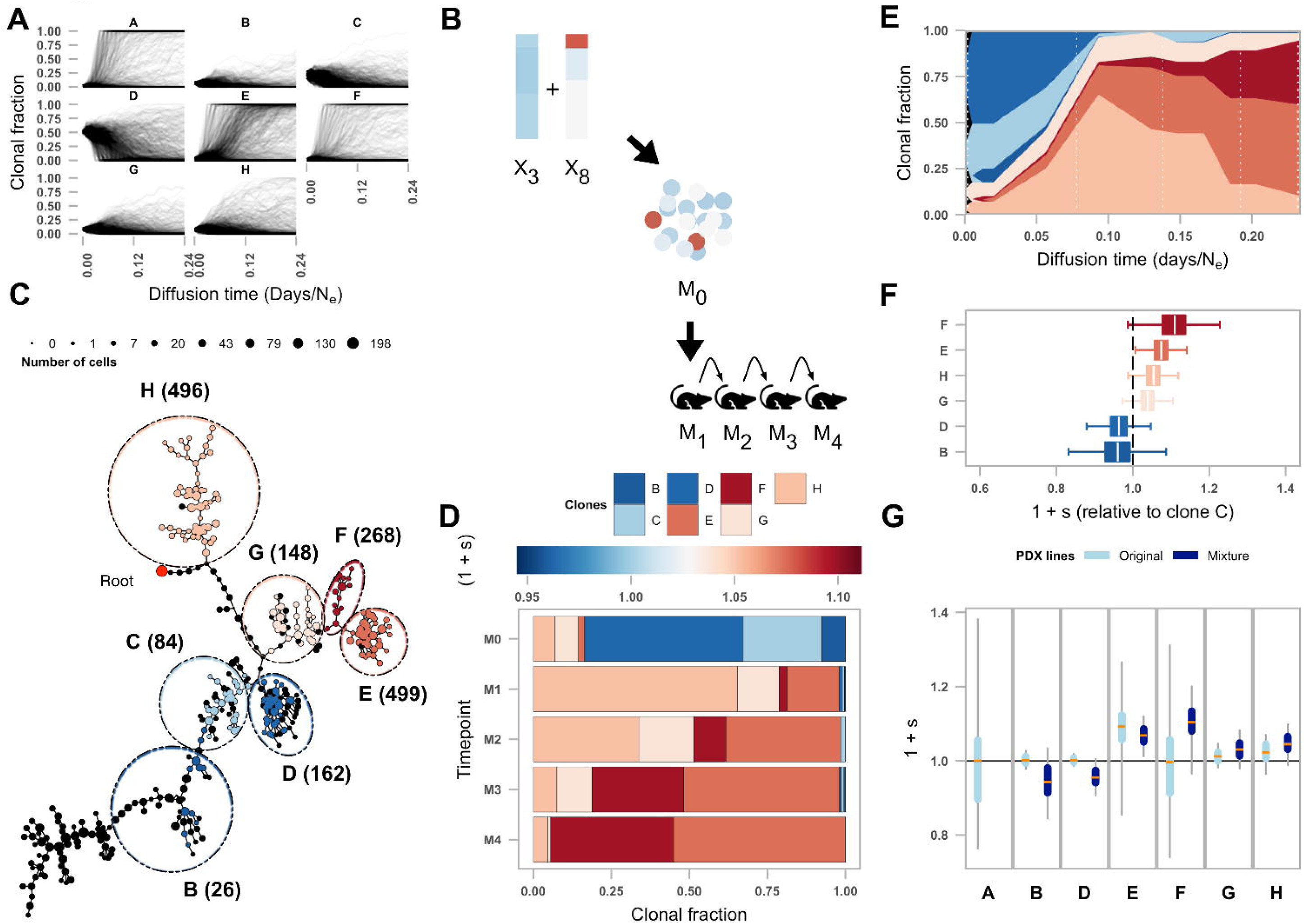
Mixture experiment on TNBC clones. **A)** Forward simulations using inferred selective coefficients and starting population proportions in the initial experimental mixture. Simulated trajectories are shown, exhibiting the probability of fixation of each clone. **B)** Clonal proportions of X3 and X8 to generate the initial mixture M0 and subsequent serial passaging, yielding 4 samples M1-M4. **C)** Phylogeny showing cells observed in the mixture timeseries. **D)** Clonal dynamics of mixture timeseries. **E)** Inferred trajectories. **F)** Selective coefficients of `fitClone` fit to M1-M4 clonal abundance observations. **G)** Comparison of selective coefficient posterior distributions in original and mixture timeseries.

### Clone-specific genotypes underpin clone-specific gene expression programs

We next profiled the impact of clone specific gene expression changes as a higher order representation of phenotypic properties. We tested if the genotypes of high fitness clones exhibited changes in their transcriptional program, with scRNAseq performed on matched aliquots of samples sequenced using DLP+ (**Supplementary Table 2**, Supplementary Information). We applied a statistical model, `clonealign`^30^, to map scRNAseq transcriptional profiles to their clone of origin, and to investigate gene dosage effects of copy number alterations on transcription. scRNAseq embeddings showed a dynamic pattern of global expression over time which tracked with clone assignments (**Fig. 6A**), indicating co-variation of transcriptional properties with clonal abundance. Pairwise comparisons of clone-specific differential gene expression over the genome revealed spatially correlated distributions reflective of gene dosage effects (**Supplementary Fig. 6**) at the level of whole chromosomes, whole arm and segmental aneuploidies. Transcriptional profiles from late timepoints (when high fitness clones had grown in abundance) indicated copy number driven gene expression in each of the *p53-/-a, p53-/-b* and TNBC series (**Fig. 6B**). DLP+ and `clonealign` clone abundance measures were positively correlated across all libraries (Pearson correlation coefficient = 0.94, p*<*0.001, **Fig. 6C**), consistent with clone specific gene dosage effects in the majority of libraries. Examples from *in vitro* and PDX series showed between 17 and 42% of all differentially expressed genes have clone specific copy number differences (**Fig. 6D**, FDR*<*0.01). In high fitness clones with high level amplifications as distinguishing features, we noted accompanying in *cis* clone-specific differential gene expression in *PVT1* and *MYC* in passage X57 of *p53-/-a* (clone G with 9 copies of Chr8q), *PFDN4* in passage X50 of *p53-/-b* (clone D with 7 copies of Chr20q) and *VCX3B* in passage X10 of TNBC (clone E with 8 copies of ChrXp) (**Fig. 6E**). Together these data indicate that clonal genotypes driving high fitness trajectories are accompanied by changes in gene expression at both chromosomal and focal level copy number alterations.

**Figure 6.**
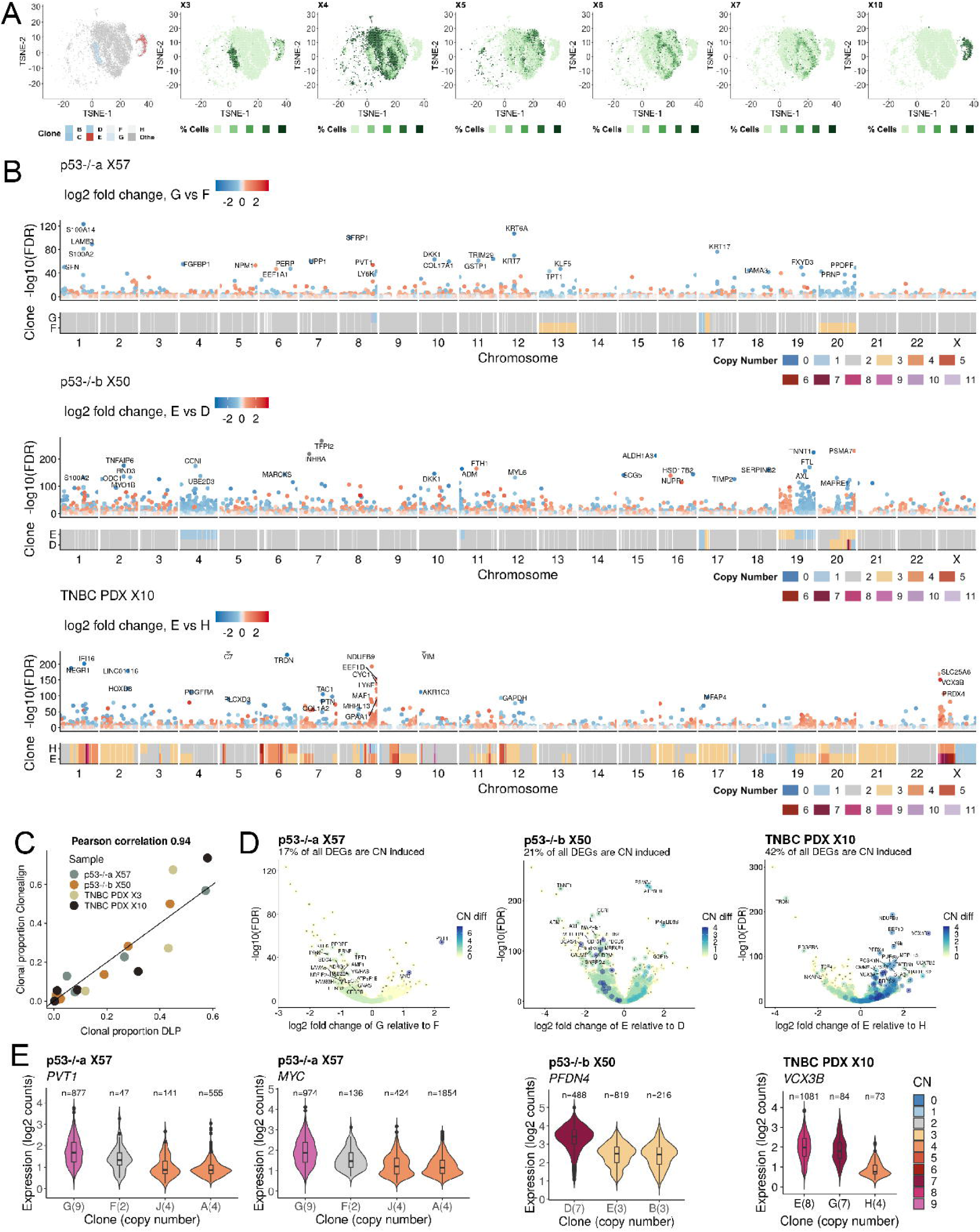
Gene expression impacts of clone-specific copy number profiles. **A)** Low dimensional t-SNE embedding of matching scRNAseq libraries across the TNBC timeseries annotated with X3 and X10 `clonealign` assignments (left) and density of cells over the timeseries X3-X10. **B)** Genome-wide view of differential gene expression between pairs of clones: G *vs* F in *p53-/-a* in sample X57 (top), E *vs* D in *p53-/-b* in sample X50 (middle), E *vs* H in TNBC sample X10 (bottom). **C)** `clonealign` clonal proportions vs DLP+ clonal proportions indicating positive correlation. **D)** Volcano plots −log10(FDR) plotted against log2 fold change of pairwise differential gene expression between clones. **E)** Distributions of clone-specific gene expression for genes in high-level amplifications in *p53-/-a, p53-/-b* and TNBC.

### Clonal competition and fitness costs of platinum resistance

We tested how pharmacologic perturbation with cisplatin impacted the stability of the fitness landscape of the TNBC series. We generated a separate branch of the TNBC model where we administered cisplatin (2mg/kg, *Q3Dx8* i.p. max) serially over four successive passages to induce drug resistance (**Supplementary Fig. 5B,C**, Supplementary Information). For each serially treated tumour, a parallel set of transplanted mice were left untreated, establishing corresponding drug ‘holiday’ samples (**Fig. 1A**). We coded the treated passages with ‘T’ and untreated with ‘U’, initialised by the X3 untreated (U) passage. The first treatment passage (*X4 UT*) exhibited rapid tumour shrinkage (*>*50% of initial size). However *X5 UTT, X6 UTTT* and *X7 UTTTT* had progressively less response, indicating drug resistance and positive growth kinetics (**Supplementary Fig. 5E**). Decomposing the growth dynamics over (*X3 U*; *X4 UT*; *X5 UTT*; *X6 UTTT*; *X7 UTTTT*) into clonal trajectories with DLP+ analysis suggested sustained cisplatin treatment inverted the fitness landscape. A new Clone R, derived from Clone A in the phylogeny, but with a distinct clonal genotype (fewer copies of *MYC* and deletions at *RB1, PRDM9* and *NUDT15* loci (**Fig. 7A,B**)), swept to fixation comprising 48% (*X4 UT*), 98% (*X5 UTT*), 100% (*X6 UTTT*) and 100% (*X7 UTTTT*) of cells across the treated series (**Fig. 7C**, **Supplementary Fig. 3**). Notably, the high fitness clones E, H, G, D from the untreated series exhibited low fitness coefficients in the treatment series and were no longer detected (**Fig. 7D**, **Supplementary Fig. 7**). Conversely, Clones A, B, and C, comprising a low fitness phylogenetic superclade, distinct from high fitness clones E and F in the untreated series, were the precursors to the resistant clone R (**Fig. 7E**). Thus, cisplatin perturbation resulted in a near complete inversion of the fitness landscape.

**Figure 7.**
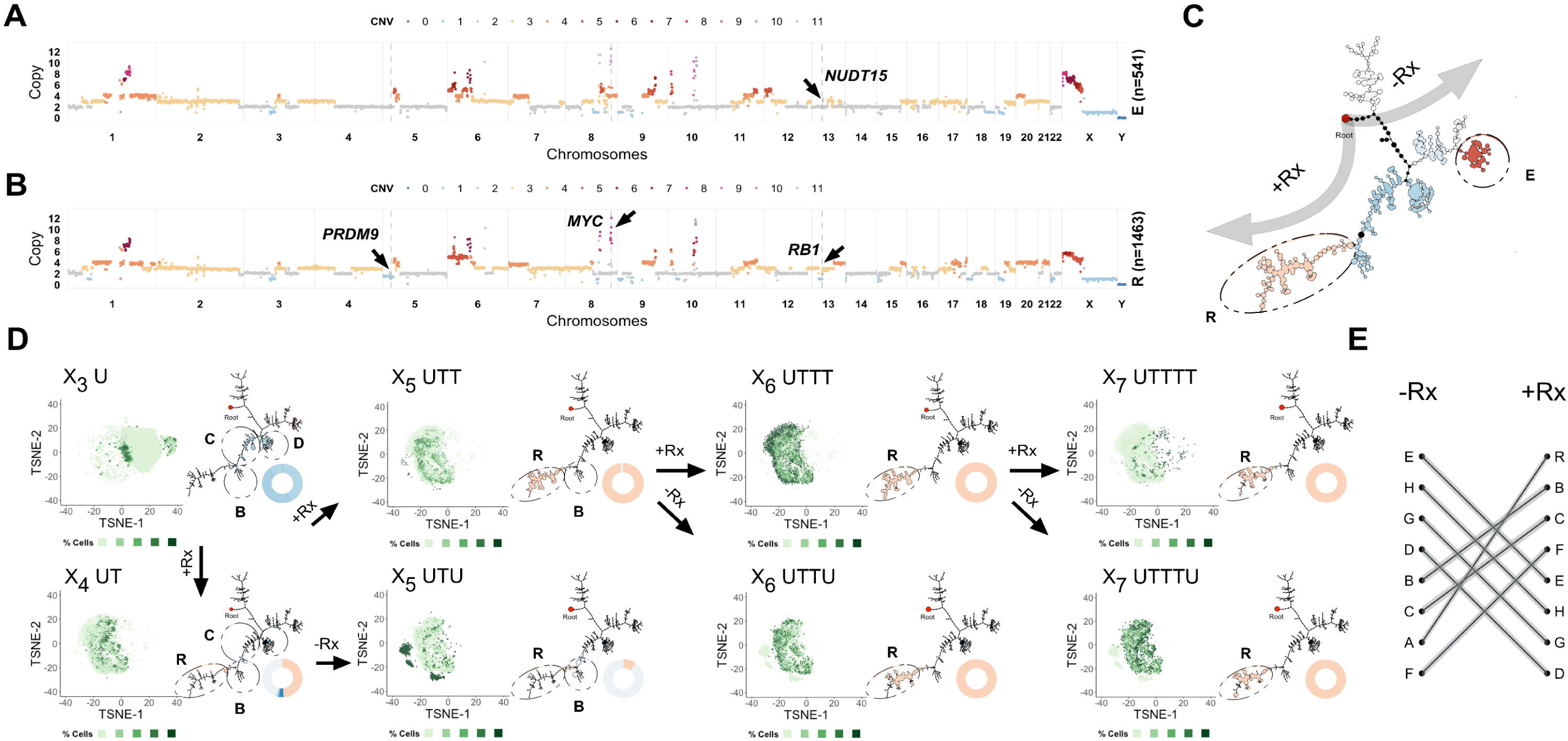
Impact of pharmacologic perturbation with cisplatin on fitness landscapes. **A)** Copy number genotype of clone E from untreated timeseries. **B)** Copy number genotype of clone R from treated timeseries (arrows indicate differences to clone E). **C)** Direction of evolutionary flux in the untreated (-Rx) and treated (+Rx) series. **D)** Evolution and transcriptional changes as a function of drug treatment and drug holiday. Arrows indicate the path of serial passaging. For each sample, the phylogeny with clonal abundance from DLP+ and the scRNAseq embedding is shown, reflecting selection and phenotypic plasticity. **E)** Inversion of the fitness landscape – note clone A maps to clone R in post treatment since R is derived from A, but only emerges in the treatment series.

We next asked whether the clonal dynamics in the presence of cisplatin were reversible by examining the drug holiday samples (**Fig. 1A**, *X5 UTU*; *X6 UTTU*; *X7 UTTTU*). In the first drug holiday *X5 UTU*, clonal composition reverted to consist predominantly of precursor clone B with 90% abundance, and only 10% abundance from clone R (**Supplementary Fig. 3**). However, in *X6 UTTU* and *X7 UTTTU* no reversion was detected, and these populations consisted of *>*99% Clone R, similar to their on-treatment analogues. Thus, clonal competition in the absence of drug led to clones derived from the A-B clade outcompeting clone R, and clone-specific cisplatin resistance thus has a fitness cost. Moreover, the genotype specificity of reversion between *X4 UT* to *X5 UTU* indicates that the clonal dynamics can be attributed to selection of genomically defined clones with differential fitness.

While transcription phenotype clusters tracked with genotypic clones in the untreated series (**Fig. 6**), analysis of scRNAseq from the parallel treated samples indicated that phenotypes were also impacted by cisplatin (Pearson correlation of textttDLP+ clones and `clonealign` scRNAseq clones = 0.99, **Supplementary Fig. 8A**). Global transcription effects generating increased phenotypic volume^31^ (**Fig. 8A**) and transcriptional velocity^32^ (**Fig. 8B** and **Supplementary Fig. 8B**) were observed as a function of time on treatment. As nearly all tumour cells were clone R from *X5 UTT* onwards in the treatment series, we attributed transcriptional changes to phenotypic plasticity (**Fig. 8C,D** and **Supplementary Fig. 8C,D**). Globally, 27 genes exhibited monotonic linear and coordinated activity as a function of time on treatment (**Fig. 8C**). Modified pathways included previously confirmed cisplatin-resistance metabolic processes such as oxidative phosphorylation ^33^, TNFA signaling via NFKB ^34–36^, E2F targets ^37^ and hypoxia ^38–41^ (**Supplementary Table 8**). From these pathways specific gene expression trajectories (**Fig. 8E**) exemplified systematic response to sustained treatment, relative to the untreated and holiday regimes. A minor component of the phenotypic volume was reversible (**Fig. 8A**) on drug holiday, indicating only modest or partial reversibility of gene expression of clone R (**Fig. 8E**, X5-X6, X6-X7, Rx and Rx-H) for some genes (*CEBP, NDUF7, MYC*).

**Figure 8.**
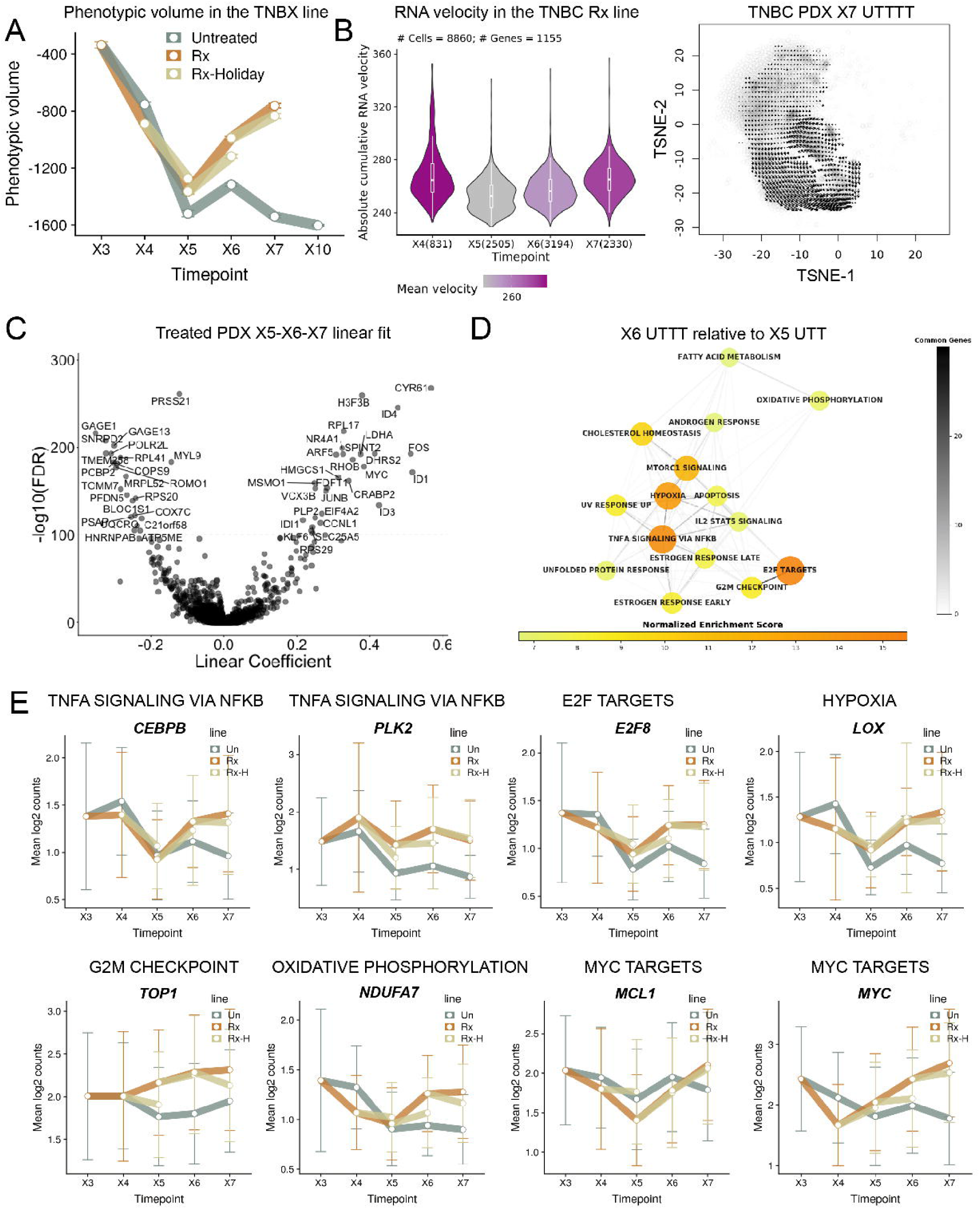
Impact of pharmacologic perturbation with cisplatin on phenotypic plasticity. **A)** Phenotypic volume as a function of time and treatment status. **B)** Transcriptional velocity for the treated timeseries: distribution of cell velocities (left) and visualization of velocity vectors at *X7 UTTTT* (right). **C)** Genes that are significantly increased (positive x axis) or decreased (negative x axis) across *X5, X6* and *X7* treated samples, obtained by fitting a linear model to the expression levels of these three samples. **D)** Pathway enrichment network comparing *X6 UTTT* against *X5 UTT*. All listed pathways are enriched at the later timepoint. **E)** Examples of gene expression values in all scRNAseq TNBC PDX libraries, confirming existing literature for cisplatin resistance.

Together, these data show the impact of cisplatin selective pressure on the starting tumour cell population is reversible while genomic clonal competition with precursor clones is still possible, but becomes fixed once the evolutionary bottleneck narrows and purifies the population. Once fixed, a minor component of the expression landscape remains reversible. Thus, cisplatin resistance occurred in phases – first dominated by clonal selection on mixed populations, followed by transcriptional plasticity on a fixed genotype.

## Discussion

Here we show that decoding the contributions of clonal competition and expression landscapes in the course of tumour growth with and without drug selection can be achieved by measuring and modeling cellular dynamics with granular timeseries over several months-years. Single cell whole genome sequencing allowed for robust application of population genetics statistical models and accompanying single cell transcriptome sequencing allowed for decoupling of clonal selection and transcriptional plasticity. *TP53* ablation in diploid mammary epithelial cells resulted in clonal expansions driven by copy number alterations, with whole genome, whole chromosome and focal alterations all leading to fitness advantages over diploid baseline. As point mutations did not show evidence of association with increased fitness, our results suggest that positive selection attributed to copy number changes of all scales may be under-appreciated. This has implications for interpreting etiologic processes of *TP53* driven cancers where the rates of structural variation acquisition and deviation away from diploid configurations confers quantitative fitness advantages. In tumour evolution within patients, copy number alterations as a key biological process is revealing impact on treatment outcomes and co-morbidities ^42^, innate and adaptive immune response ^43, 44^, the root cause of major genomic reconfigurations ^45^ and evolutionary plasticity ^13^. Our work here suggests that *TP53* mutation can lead to structural alterations *in vitro* that are nevertheless observed in breast cancer patients, thereby representing a realistic model for studying how the impact of driver mutations inducing genomic instability leads to clonal expansions and evolutionary selection. Variously through single cell approaches ^46^ or computational reconstruction of evolutionary histories ^6^, the evolutionary impact of structural variations is still a work in progress. Over successive generations *in vitro* and in PDX, in the context of pharmacologic and mutational perturbation, emergent copy number changes contribute to the kinetics of the fitness landscape, consistent with a continual diversifying mechanism that induces competitive clonal advantages.

The impact of drug intervention on cancer evolution is a key determinant of patient outcomes across all human cancers. We show that resistance to platinum, a commonly used chemotherapeutic agent, can be multi-faceted, involving time dependent phases of clonal selection pruning unfit clones and phenotypic plasticity regulating global transcription with metabolic and cell-cycle pathways. As clonal drug resistance was consistent with a fitness cost, we suggest this could be exploited in future therapeutic strategies. Forecasting the trajectories of cancer clones is of immediate importance to understanding therapeutic response in cancer and for deploying adaptive approaches ^3^. The presence within a tumour of lineage precursors to resistant genotypes may define time windows within which clonal competition could mediate plasticity to treatment. We suggest that population genetics modeling of timeseries tumour or tumour cell-free DNA measurements to predict clonal evolution is tractable, but will require complementary measurements of genotypic clonal abundance and phenotypic plasticity to gain comprehensive understanding. Further study with timeseries modeling will provide insight into therapeutic strategies promoting early intervention, drug combinations and evolution-aware approaches to clinical management ^47^.

## Methods

All methods are detailed in the Supplementary Information. Software and source code implementing analytical and statistical methods available on github: [https://github.com/UBC-Stat-ML/fitclone]. Raw sequencing data for DLP+ and 10x scRNASeq will be available from the European Genome-Phenome archive prior to publication.

## Author contributions

SPS and SA: project conception and oversight, manuscript writing, senior responsible authors; ABC: statistical inference method development and oversight; SS: computational method development, data analysis, manuscript writing; FK: mouse modeling, tissue procurement, data generation, manuscript writing; NC, MA, MJW, KC, AZ, FD, JP, DG, DL, AM: computational biology, data analysis; COF, TM, BW, JB, JB, JT, EL, HL, TA, SL, BYCC, PE, TK: tissue procurement, biological substrates and data generation; RM, AJM, MAM: genome sequencing; NR: manuscript editing

## Acknowledgements

This project was generously supported by the BC Cancer Foundation at BC Cancer and Cycle for Survival supporting Memorial Sloan Kettering Cancer Center. SPS holds the Nicholls Biondi Chair in Computational Oncology and is a Susan G. Komen Scholar. SA holds the Nan and Lorraine Robertson Chair in Breast Cancer and the Canada Research Chair in Molecular Oncology and CRC Chair in Molecular Oncology. Additional funding provided by Cancer Research UK Grand Challenge Program, the Canadian Cancer Society Research Institute Impact program and Terry Fox Program Project Grant awards to SA, SPS and ABC.

## Competing Interests

SPS and SA are shareholders and consultants of Contextual Genomics Inc.

