## Supplementary information, supplementary methods, and supplementary tables for "Single cell fitness landscapes induced by genetic and pharmacologic perturbations in cancer"

1. Department of Molecular Oncology, BC Cancer, Vancouver, BC, Canada
2. Department of Pathology and Laboratory Medicine, University of British Columbia, Vancouver, BC, Canada
3. Computational Oncology, Department of Epidemiology and Biostatistics, Memorial Sloan Kettering Cancer Center, New York, NY 10065, USA
4. Department of Computer Science, University of British Columbia, Vancouver, BC, Canada
5. Department of Statistics, University of British Columbia, Vancouver, BC, Canada
6. Lunenfeld-Tanenbaum Research Institute Mount Sinai Hospital Joseph & Wolf Lebovic Health Complex, Molecular Genetics, University of Toronto, Toronto, ON, Canada
7. Canada's Michael Smith Genome Sciences Centre, BC Cancer, Vancouver, BC, Canada
8. CRUK Grand Challenge IMAXT Team

* - equal contribution

† - Corresponding Authors:



**Keywords:** tumour evolution, single cell sequencing, fitness, timeseries, phylogenetic reconstruction, drug resistance

### 1 Human mammary cell lines and serial passaging

The human mammary epithelial cell line wild type 184-hTERT and isogenic 184-hTERT-P53 KO cell line, generated from 184hTERT WT-L9, were grown as previously described [1, 2]. Two branches of 184-hTERT-P53$^{-/-}$ (clone 95.22) along with the counterpart wild type branch were serially passaged for $\approx$ 55-60 passages, by seeding $\approx$ 1 million cells into a new $10\,\mathrm{cm}$ tissue culture treated dish (Falcon- CABD353003) and cryopreserving every fifth passage, using the growth media, MEGM™Mammary Epithelial Cell SingleQuot Kit Supplements and Growth Factors (Lonza CC-4136), with $5\,\mathrm{\mu g\,ml^{-1}}$ transferrin (Sigma) and $2.5\,\mathrm{\mu g\,ml^{-1}}$ isoproterenol (Sigma) as previously described [2]. Cells were grown to around 85-90% confluence, trypsinized for 2 minutes (Trypsin/EDTA 0.25%(VWR CA45000-664)), re-suspended in cryopreservation medium (10% DMSO-Sigma-D2650, 40% FBS-GE Healthcare SH30088.03, 50%media) and frozen to $-80\,^{\circ}\mathrm{C}$ at a rate of$-1\,^{\circ}\mathrm{C\,min^{-1}}$. Cells were cultured continuously from passage 10 (post initial cloning as in [2]) to passage 60 for 184-hTERT WT and upto passage 57 and passage 55 for the P53$^{-/-}$ branches a and b respectively, from initial cloning/isolation (**Supplementary Fig. 2**A. Genome sequencing was undertaken at passages 25,30, 51 and 60 from the wild type branch, passages 10,15,25,30,40,50, and 57 from P53 KO branch a and passages 20, 30, 35, 40, 45, 50 and 55 from P53 KO branch b. Also, the transcriptome sequencing was carried out on passages 11 and 57 of P53$^{-/-}$ branch a and passages 15, 30 and 50 of P53$^{-/-}$ branch b (see Methods). The TP53 alleles were confirmed by Sanger sequencing as NM_000546(TP53):c.[156delA];[156delA], p.(GIn52Hisfs*71), and the absence of TP53 protein was confirmed with western blot (**Supplementary Fig. 2**B).

### 2 Establishment and serial passaging of patient derived xenografts

The Ethics Committees at the University of British Columbia approved all the experiments using human resources. Patients in Vancouver, British Columbia were recruited, and samples were collected under the tumour tissue repository (TTR-H06-00289) protocol and transplanted in mice under the Neoadjuvant PDX (University of British Columbia BC Cancer Research Ethics Board H20-00170) protocols.

After informed consent, tumour fragments from patients undergoing excision or diagnostic core biopsy, were collected. Tumour materials were processed as described in [3] and transplanted in mice under the animal resource centre (ARC) bioethics protocol (A19-0298-A001) approved by the animal care committee. Briefly, tumour fragments were chopped finely with scalpels and mechanically disaggregated for one minute using a Stomacher 80 Biomaster (Seward Limited, Worthing, UK) in $1\,\mathrm{ml}$ to $2\,\mathrm{ml}$ cold DMEM/F-12 with Glucose, L-Glutamine and HEPES (Lonza 12-719F). An aliquot of $200\,\mathrm{\mu l}$ of medium (containing cells/organoids) from the resulting suspension was used equally for 4 transplantations in mice.

Tumours were transplanted in mice as previously described [3] in accordance with SOP BCCRC 009. Briefly, female immuno-compromised, NOD/SCID/IL2r$\gamma^{-/-}$ (NSG) and NOD/Rag1$^{-/-}$Il2r$\gamma^{-/-}$ (NRG)[4] mice were bred and housed at the Animal Resource Centre (ARC) at the British Columbia (BC) Cancer Research Centre . For subcutaneous transplants, mechanically disaggregated cells and clumps of cells were re-suspended in $150\,\mathrm{\mu l}$ to $200\,\mathrm{\mu l}$ of a 1:1 v/v mixture of cold DMEM/F12: Matrigel (BD Biosciences, San Jose, CA, USA). 8-12 weeks old mice were anesthetized with isofluorane, then the mechanically disaggregated cells/clumps suspension was injected under the skin on the left flank using a $1\,\mathrm{ml}$ syringe and 21gauge needle. The animal care committee and animal welfare and ethical review committee, the University of British Columbia (UBC), approved all experimental procedures.

**2.1 Histopathology of PDX tumours** The hormone receptor status of both tumour samples was determined by immunohistochemistry and FISH (Fluorescence in situ hybridization) copy number. Two separate tissue microarrays were prepared using duplicate $1\,\mathrm{mm}$ cores extracted from formalin-fixed paraffin-embedded blocks containing material from passage 1 to passage 10 of both patient derived xenografts (TNBC, HER2+) used for this study. Deparaffinized $4\,\mathrm{\mu m}$ sections of paraformaldehyde fixed tumours were processed for immunohistochemistry (IHC) using a Discovery XT automated system (Ventana Medical Systems, Tucson, AZ, USA). EGFR, INPP4B, Ki67, PR, ECAD were all performed on the Ventana Discovery XT platform using CC1 for antigen retrieval, incubating for one hour at room temp, and using the UltraMap DAB detection kit. Primary antibodies to ER$\alpha$ (clone SP1, Ventana), HER2 (clone 4B5, Ventana), EGFR (clone EP22, Epitomics, Burlingamen, CA, USA) and Ki67 (clone SP6, Thermo Scientific) (**Supplementary Table 6**). Horseradish peroxidase-conjugated Discovery Universal Secondary Antibody (Ventana) was then applied and the slides developed using 3,3'-diaminobenzidine Map Kit (Ventana). The slides were reviewed by a pathologist.(**Supplementary Fig. 2**E, **Supplementary Table 7**). H&E and immunohistochemistry (IHC) of the two PDX tumours showed that TNBC is derived from a triple negative breast cancer patient while HER2+ was derived from a HER2+ breast cancer patient. HER2 IHC was scored as 2+, HER2/CEP17 ratio was calculated as 6.5 (positive)[5] from FISH and focal high level amplification (average copy number state of 10) of the *ERBB2* locus (approximately Chr17:37500001-38000000) was found in the DLP+ data.

**2.2 Serial passaging of PDX** Tumours were serially passaged as described [3]. Briefly, for serial passaging of PDX, xenograft-bearing mice were euthanized when the size of the tumours approached $1000\,\mathrm{mm}^3$ in volume (combining together the sizes of individual tumours when more than one was present). The tumour material was excised aseptically, then processed as described for primary tumour. Briefly, the tumour was harvested and minced finely with scalpels then mechanically disaggregated for one minute using a Stomacher 80 Biomaster (Seward Limited, Worthing, UK) in $1\,\mathrm{ml}$ to $2\,\mathrm{ml}$ cold DMEM-F12 medium with Glucose, L-Glutamine and HEPES. Aliquots from the resulting suspension of cells and fragments were used for xenotransplants in next generation of mice and cryopreserved. Serially transplanted aliquots represented approximately 0.1-0.3% of the original tumour volume. HER+ and TNBC PDX were passaged upto 10 generations and scWGS was carried out at each timepoint while scRNAseq was done at initial, mid and late timepoint(**Supplementary Fig. 2**C).

**2.3 TNBC PDX timeseries treatment with cisplatin** NRG mice of the same age and genotype as above were used for transplantation treatment experiments. Drug treatment with cisplatin (Platinum) was commenced when the tumour size reached approximately $300\,\mathrm{mm}^3$ to $400\,\mathrm{mm}^3$. Cisplatin (Accord DIN: 02355183) was administered i.p. at $2\,\mathrm{mg\,kg}^{-1}$ every third day for 8 doses maximum (Q3Dx8). The dosage schedule was adjusted 50% less than what is mentioned in the literature [6, 7] and around one third of the maximum tolerated dose (MTD) calculated in the immunodeficient mice (**Supplementary Fig. 5**B). Low dose cisplatin pulse and tumour collection timings were optimized to achieve the experimental aims of tumour resistance. The aim was to collect tumour at 50% shrinkage (from the starting tumour at the time treatment started) in size when measured with a caliper. Cisplatin $1\,\mathrm{mg\,ml}^{-1}$ was diluted in 0.9% NaCl to obtain concentrations $200\,\mu\mathrm{l}/20\,\mathrm{g}$ of mouse weight and kept in glass vials at room temperature. Quality control (QC) samples were prepared freshly on each day prior to the dosing. Mice were continually monitored for acute signs of toxicity including pain at injection site, skin tenting, coat scruffing, sunken eyes, food consumption and behaviour for the first two hours following compound administration. For TNBC PDX, 8 mice at passage four (X4) of were transplanted in parallel for the treatment/treatment holiday study group. Half of the mice were treated with cisplatin when tumours exhibited $\approx 50\%$ shrinkage, the residual tumour was harvested as above and re-transplanted for the next passage (X5) in the group of eight mice. Again, half of the mice at X5 were kept untreated while the other half were exposed to cisplatin following the same dosing strategy. Four cycles of cisplatin treatment were generated, with a parallel drug holiday group at each passage. Cisplatin treated tumours were labelled as *UT,UTT,UTTT,UTTTT* for each of the four cycles of drug respectively, while the tumours on drug holiday were called as *UTU, UTTU* and *UTTTU* for the three timepoints. ScWGS and scRNAseq was carried out from each tumour during the timeseries treatment with counterpart drug holiday and untreated controls. (**Supplementary Fig. 5**C).

**2.4 PDX tumour growth measurement curves** NRG mice received sub-cutaneous inoculation (SQ) of tumour cells ($150\,\mu\mathrm{l}$) on day 0. The tumours were allowed to grow to palpable solid nodules. Around 7-9 days after they are palpable, their size were measured with calipers every 3rd day. tumours were measured in two dimensions using a digital caliper and expressed as tumour volume in mm3; defined as: [volume= $0.52\times$(Length)$\times$(Width)]. Both patient derived xenografts, HER2+ and TNBC exhibited progressively higher tumour growth rates in later passages (**Supplementary Fig. 5**D). Under drug perturbation, the treated tumours in the first two cycles of treatment showed rapid growth reduction but in third cycle started showing non-responsive behaviour leading to total resistance in fourth cycle (**Supplementary Fig. 5**E).

**2.5 TNBC PDX tumour mixing experiments** Frozen untreated passages three (X3) and eight (X8) vials from TNBC PDX, were thawed and remixed in two different volumetric proportions of X3:X8 by tumour weight (1:1 and 1:0.25-data from 1:0.25 not shown). From each of different dilutions, $200\,\mu\mathrm{l}$ of aliquot was taken to be used to transplant in two mice each using the same protocol of transplantation as described above. Before transplantation a small proportion of the physical mixture of cells, from the 1:1 ratio, was subjected to whole genome single cell sequencing to measure the baseline clonal composition labelled as M0 and its subsequent PDX as M1 (**Supplementary Fig. 5**A). Each of the thawed X3 and X8 cell populations used for mixing were also transplanted independently to confirm the viability of the tumour material for PDX tumour growth. The tumour cell mixture was then serially passaged over 4 generations, designating the transplants as M1-M4. Tumours from each X3:X8 serial passage were collected and analysed with scWGS (DLP+) as for other samples.

### 3 Single cell whole genome sequencing and library construction with DLP+

All libraries, including metrics on number of cells, average number of reads per cell and quality control metrics are listed in **Supplementary Table 1**.

**3.1 Creation of single cell suspensions from PDX** Tumour fragments from PDX samples were incubated with a collagenase/hyaluronidase 1:10 (10X) enzyme mix (STEM CELL technologies, Catalog #07912) in 5 ml DMEM/F-12 with Glucose, L-Glutamine and HEPES (Lonza 12-719F)and 1%BSA at 37 °C with intermittent gentle pipetting up and down the sample every 30 min for 1 min, during the first hour with a wide bore pipette tip, and every 15-20 min for the second hour, followed by centrifugation (1100 rpm, 5 min) and supernatant removal. The tissue pellet was resuspended in 1 ml of 0.25 percent trypsin-EDTA (VWR CA45000-664) for 1 min, superadded by 1 ml of DNAse 1/dispase 100 µl/900 µl (StemCell 07900,00082462) pipetted up and down 2 min, followed by neutralization with 2% FBS in HBSS with 10 mM HEPES (STEMcells Catalog #37150). This cell suspension was then passed through a 70 µm filter to remove remaining undigested tissue and centrifuged for 5 min at 1100 rpm after topping it up to 5 ml with HBSS. Single cells pellet was resuspended in PBS + 0.04% BSA (Sigma) in appropriate volume to achieve ≈ 1 million per ml concentration of cells for robot spotting for DLP+.

**3.2 Robot spotting of single cells into the nanolitre wells and library construction** scWGS DLP+ library construction was carried out as described in [1]. Briefly, single cell suspensions from cell lines and patient derived xenografts were fluorescently stained using CellTrace CFSE (Life Technologies) and LIVE/DEAD Fixable Red Dead Cell Stain (ThermoFisher) in a PBS solution containing 0.04% BSA (Miltenyi Biotec 130-091-376) incubated at 37 °C for 20 minutes. Cells were subsequently centrifuged to remove stain, and resuspended in fresh PBS with 0.04 percent BSA. This single cell suspension was loaded into a contactless piezoelectric dispenser (sciFLEXARRAYER S3, Scienion) and spotted into the open nanowell arrays (SmartChip, TakaraBio) preprinted with unique dual index sequencing primer pairs. Occupancy and cell state were confirmed by fluorescent imaging and wells were selected for single cell copy number profiling using the DLP+ method [1]. Briefly, cell dispensing was followed by enzymatic and heat lysis. After cell lysis, tagmentation mix (14.335 nL TD Buffer, 3.5 nL TDE1, and 0.165 nL 10% Tween-20) in PCR water were dispensed into each well followed by incubation and neutralization. Final recovery and purification of single cell libraries was done after 8 cycles of PCR. Cleaned up pooled single-cell libraries were analyzed using the Aglient Bioanalyzer 2100 HS kit. Libraries were sequenced at UBC Biomedical Research Centre (BRC) in Vancouver, British Columbia on the Illumina NextSeq 550 (mid- or high-output, paired-end 150-bp reads), or at the GSC on Illumina HiSeq2500 (paired-end 125-bp reads) and Illumina HiSeqX (paired-end 150-bp reads). The data was then processed to quantification and statistical analysis pipeline [1].

### 4 Single cell RNA sequencing (scRNAseq)

All libraries generated using 10x scRNAseq are listed in **Supplementary Table 2**.

**4.1 Processing of cell lines for scRNAseq data** Suspensions of 184-hTERT *p53 WT* and KO cells were fixed with 100% ice-cold methanol prior to preparation for scRNAseq. Single cell suspensions were loaded onto the 10x Genomics single cell controller and libraries prepared according to the Chromium Single Cell 3' Reagent Chemistry kit standard protocol. Libraries were then sequenced on an Illumina Nextseq500/550 with 42bp paired-end reads, or a HiSeq2500 v4 with 125bp paired-end reads. 10x Genomics Cell Ranger 3.0 was used to perform demultiplexing, alignment and counting.

**4.2 Processing of patient derived xenografts for scRNAseq data** PDX tumours were harvested and mechanically disaggregated into small fragments to viably freeze them according to the protocol mentioned above. One of the viable frozen tumour vial was thawed and after washing out the freezing media, the tumour clumps and fragments were incubated with digestion enzymes as with DLP+ preparation. After complete dissociation with collagenase/hyaluronidase enzyme mix according to the protocol at 37 °C, followed by briefly washing in 0.05% trypsin-EDTA and resuspension in 0.04% BSA in PBS. Dead cells were removed using the Miltenyi MACS Dead Cell Removal kit and cells were processed as previously described [8]. To avoid processing artifacts, dissociation methods and times were tightly controlled and for treatment and treatment holiday pairs, library construction was performed on the same chips. Library construction sample batch groupings are listed in **Supplementary Table 2**.

### 5 DLP+ sequence analysis, copy number determination and quality control filtering

FASTQ pre-processing, sequence alignment, quality control, copy number calling and S-phase classification and filtering was performed on all libraries as detailed in [1]. Briefly, cells were assigned a QS score for data quality based on a 13 feature Random forest classifier fitted and applied as per [1]. Copy number alterations on a per cell basis were determined using a hidden Markov Model (HMM) approach using the `HMMCopy` package with parameterizations detailed in [1]. S-phase cells were identified using an automated classifier trained using cell-cycle sorted cells and where features from HMM output were used to identify cells most probably in early or late phase replication of their genomes. As these cells interfere with downstream phylogenetic analysis, they were removed from the analysis according to parameter settings and thresholds established in [1]. To further enable phylogenetic inference, 10-15% of cells with highest average copy number (CN) state jumps were removed. Upon inspection, these cells included early and late dividing cells that were not captured by the s-phase classifier. Attrition of cells at each stage of quality control is shown in **Supplementary Table 1**.

**5.1 Phylogenetic tree inference, clone determination and clonal abundance measurements** We developed a single cell Bayesian tree reconstruction method based on copy number change point binary variables called `sitka` [9] to fit phylogenetic trees to the copy number profiles. In the output of `sitka`, cells are the terminal leaf nodes of the phylogenetic topology. The inferred trees were post-processed to identify clonal populations from major clades. With clonal populations defined, their abundances were counted as a function of timeseries and these were used for fitness inference (see below). Clones were constructed by identifying connected components (each a clade or a paraphyly) in the phylogenetic tree reconstruction. The tree was 'cut' into discrete populations according to the following procedure (see Algorithm 1). Let $L$ be a set of loci and $C$ a set of cells and $|L|$. Define $\tau = (L, C, E)$ to be a rooted phylogenetic tree with $E$ its set of directed edges. The phylogenetic tree is assumed to comprise internal nodes that are phylogenetic markers (loci) and terminal nodes that are either cells or loci. Terminal loci are considered unused phylogenetic markers and are discarded. Let $|\tau| = |C|$, that is the number of cells that belong to tree $\tau$. By $\tau_l = (L_l, C_l, E_l)$ denote the subtree rooted at node $l$. Set pa($l$) and child($l$) be the parent node and set of immediate children of node $l$ with desc($l$) comprising all its descendant.

The inputs to the algorithm are the rooted phylogenetic tree $\tau$ and the copy number states of its cells. A clone is defined as connected components (each a clade or a paraphyly) in the graph tree $\tau$ composed of cell of sufficient genomic homogeneity. The degree of homogeneity can be tuned by limiting the number of loci and the difference in copy number of sub-clades in a clone. The algorithm works by first finding the coarse structure, that is dividing the tree into major clades and then looking for fine structures within each clade by traversing the tree in a bottom up manner and merging loci that are sufficiently similar. The remaining loci constitute the roots of detected clades.

To obtain the coarse structure from the reconstructed phylogenetic tree we use a two step procedure. (i) First we identify monophyletic clades via 1. (ii) We then remove the cells comprising the clades found in step one from the tree and repeat algorithm 1. We note that these new clades (if any) could be paraphyletic.

To find the fine structures within the initial clades we use the following procedure. For each clade and its corresponding sub-tree $\tau_s$, denote by $L_s$ a set of loci $l$ for which $m \leq |\tau_l| \leq M$. In a bottom-up traverse of the tree, for each node $l \in L_s$, remove it from $L_s$ if $|\tau_{\text{pa}(l)}| - |\tau_l| \leq n$, otherwise remove $\tau_l$ from $\tau_c$ At the end of the tree traversal set $L_s$ contains new candidate roots for each initial clade. For each $l \in L_s$ define the summary copy number profile as a vector whose $i$th element is the median of the copy number states of the $i$th for all cells in $\tau_l$. Compute the distance between two subclades as the mean absolute difference of their median genotypes. Merge subclones induced by $L_s$ if their summary copy number profiles are too similar. We can do this by computing a t-test over the pairwise distances to exclude outlier subclades and merge the rest. For the cell lines datasets, namely *p53 WT* and *p53-/-a* and *p53-/-b*, we opted to also split clades by the ploidy of their constituent cells, where ploidy is defined as the most recurrent CN state in the cell.

Once clones are identified, we set the abundance of each clone at a specific timepoint as the fraction of cells in that clone from that timepoint. We note that for the data from WGS bulk sequencing [3] we used the following procedure to estimate clonal fractions: (i) let $M$ denote the mutational cellular prevalence (rows) estimated over multiple timepoints (columns) using the multi-sample `PyClone` [10] model, (ii) define $A$ as the genotype matrix (which mutation-cluster (rows) is present in which clones (columns), (iii) then we set $AX = M$ where $X = A^{-1}M$ are the clonal fractions overtime. (iv) we solve for $X$ using QR-decomposition.

**Algorithm 1** Heuristic (Top-down)

**procedure** SITKA-LUMBERJACK($\tau$)
    locus-queue $\leftarrow$ DepthFirstSearch($\tau$)
    **for** locus $l$ in locus-queue **do**
        **if** $m \leq |\tau_l| \leq M$ **then**
            $CUTS \leftarrow CUTS \cup l$
            Remove all loci below $l$ from $\tau$
        **else**
            **for** locus $l'$ in descendants(l) **do**
                **if** $m \leq |\tau_{l'}| \leq M$ **then**
                    $CUTS \leftarrow CUTS \cup l'$
                    Remove all loci below $l'$ from $\tau$
            **if** no eligible loci were found **then**
                $CUTS \leftarrow CUTS \cup l$
                Remove $\tau_l$ from $\tau$
    **return** $CUTS$

**5.2 Single nucleotide variant analysis from DLP+** A group of SNV loci candidates was selected from the aggregated data in which all cells were collapsed into a pseudo-bulk genome. Host mouse infiltrating stromal cells were filtered from the PDX libraries based on proportion of reads aligning to the mouse genome. Mouse cells were filtered from xenograft libraries using fastqscreen (`https://www.ncbi.nlm.nih.gov/pubmed/30254741.2`). Fastqscreen was used to determine, for each read, whether that read originated from human, mouse and salmon DNA. Cells for which 5% or more reads mapped exclusively to either mouse or salmon were determined to be non-human or doublets and were excluded from further analysis. Attrition of cells after quality control is shown in **Supplementary Table 1**. MutationSeq and Strelka were run for selecting SNV loci candidates from the pseudo-bulk sample [11, 12] as per [1]. The loci with MutationSeq probability greater than 0.9 and Strelka's Phred quality score of 20 or more were identified as SNV candidates. In addition, each locus should demonstrate at least two variant reads across the cells. **Supplementary Table 5** shows the number of SNV candidate loci in each sample.

The `sitka` SNV calling model [9] was used to detect SNVs in each locus per each cell. This SNV calling method is a Bayesian approach that incorporates the copy number and the underlying tree to identify the mutation status of each cell. The SNV call probabilities are inferred based on the placement of SNVs in on tree. **Supplementary Fig. 3** demonstrates the mutation status of 200 random loci across each cell of the tree in cell lines and PDX libraries. The mutation calls corroborate the clone cuts of the tree while defining finer resolution.

**5.3 Mutation rate analysis** To assess the rate of mutation accumulation across time and within clones we counted the number of SNVs identified per cell as well as the number of copy number breakpoints (i.e., segments of different copy number). The number of copy number breakpoints was defined as the number of intra-chromosomal change-points identified by the hidden Markov model copy number calling algorithm. In order to assess how mutations accumulate over time, passage numbers were converted to generations using measured doubling times of the cell lines and PDX models. To statistically assess mutation accumulation over time we performed a linear regression using individual cell level mutation counts and covariates: mutation $\sim$ generation + coverage_breadth + cell_ploidy using the lm() function in R. Similarly, to evaluate which clones had the largest number of mutations we performed the following regressions mutation $\sim$ clone + coverage_breath + cell_ploidy. coverage_breadth is the proportion of the genome that was sequenced.

### 6 Fitness modelling

We describe in this section two approaches to fitness modelling. The first is a Bayesian state-space model (`fitClone`) based on the Wright-Fisher diffusion with selection (Section 6.1). The second is a deterministic logistic growth model (Section 6.9). We compare the two methods in Section 6.10. The comparison favours the Bayesian model, hence this is the model used in our results unless we specify otherwise.

**6.1 fitClone: a Bayesian fitness model for timeseries data** We developed a Bayesian model and associated inference algorithm based on a diffusion approximation to $K$-allele Wright-Fisher model with selection. We start with timeseries clonal abundance measurements over a fixed number of clones and estimate two key unknown parameters of interest: *fitness coefficients* $s_i$ for clone $i$ which represents a quantitative measure of the growth potential of a given clone; and *distributions over continuous-time trajectories*, a latent (unobserved) population structure trajectory in 'generational' time.

After briefly reviewing and setting notation for Wright-Fisher diffusions with selection (Section 6.2), we introduce the Bayesian model we used to infer quantitative fitness of clones from timeseries data (Section 6.3). We then describe our posterior inference method (Section 6.4) and ancillary methods for effective population size estimation (Section 6.5), and reference clone selection (Section 6.7).

See [13] for more background on the Wright-Fisher model and [14, 15, 16, 17, 18, 19, 20, 21, 22] for previous work on inference algorithms for Wright-Fisher models.

**6.2 Wright-Fisher diffusions with selection** Let $K$ denote the number of clones obtained using the tree cutting procedure described in Section 5.1, and denote by $Z_t = (Z_t^1, \ldots, Z_t^K)$ the relative abundance of each of the $K$ clones at time $t$ in the population. The process $Z_t$ satisfies, for all $t$, the constraints $\sum_{i=1}^K Z_t^i = 1$ and $Z_t^i \geq 0$ for $i \in \{1, \ldots, K\}$. We would like to model the process $Z_t$ using a Wright-Fisher diffusion with selection.

A Wright-Fisher diffusion can be written in stochastic calculus notation as

$$dZ_t = \mu^{s,N_\text{e}}(Z_t)\text{d}t + \sigma(Z_t)\text{d}W_t \tag{1}$$

where $\{W_t\}$ is a $K$-dimensional Brownian motion, and the functions $\mu$ and $\sigma$, defined below, respectively control the deterministic and stochastic aspects of the dynamics. For $z = (z^1, z^2, \ldots, z^K)$, the vector-valued function $\mu^{s,N_\text{e}} : \mathbb{R}^K \to \mathbb{R}^K$ is defined as

$$\begin{aligned} \mu^{s,N_\text{e}}(z) &= (\mu_1^{s,N_\text{e}}(z), \ldots, \mu_K^{s,N_\text{e}}(z)) \\ \mu_i^{s,N_\text{e}}(z) &= N_\text{e} z^i (s_i - \langle s, z \rangle), \end{aligned}$$

where $\langle x, y \rangle$ is the inner product of vectors $x$ and $y$, $N_\text{e}$, the *effective population size*, discussed in more details in Section 6.5, and the parameters $s = (s_1, s_2, \ldots, s_K)$ are called *fitness coefficients*. The interpretation of the fitness parameters is that if $s_i > s_j$, then subpopulation $i$ has higher growth potential compared to subpopulation $j$. The matrix-valued function $\sigma : \mathbb{R}^K \to \mathbb{R}^{K^2}$ is defined as

$$\begin{aligned} \sigma^2(z) &= [\sigma_{i,j}^2(z)]_{i,j \in \{1,\ldots,K\}} \\ \sigma_{i,j}^2(z) &= z^i(\delta_{i,j} - z^j) \end{aligned}$$

where $\delta_{i,j}$ is the Kronecker delta. Given an initial value $z$, we denote the marginal distribution of the process at time $t$ by

$Z_t \sim \text{WF}(s, N_\text{e}, t, z)$.

**6.3 The fitClone model** Given as input timeseries data measuring the relative abundances of $K$ populations at a finite number of timepoints, the output of the `fitClone` model is a posterior distribution over the unknown parameters of interest: the fitness parameters $s$ described in the previous section, and the continuous-time trajectories interpolating and extrapolating the discrete set of observations.

To do this, `fitClone` places a prior on the fitness parameters $s$, and uses a state space model in which the latent Markov chain is distributed according to a Wright-Fisher diffusion, and the observation model encodes noisy sampling from the population at a discrete set of timepoints.

Each component of the fitness parameter, now a random variable $S_i$, is endowed with a uniform prior over a prior range $I$,

$$S_k \sim \text{Uniform}(I), k > 1,$$

where we set $S_1 = 0$ to make the model identifiable (see Section 6.7 for details). We used $I = (-10, 10)$ in our experiments. Note that the posterior is contained far from the boundaries of this prior range in all experiments.

The initial distribution, i.e. the distribution of the value of the process at time zero, is endowed a Dirichlet distribution with hyper-parameter $(1, 1, \ldots, 1)$,

$$Z_0 \sim \text{Dirichlet}(1, 1, \ldots, 1).$$

This can equivalently be seen as a uniform distribution over the $K$-simplex.

Let $t_1 < t_2 < \cdots < t_{T-1} < t_T$ denote a set of process times at which measurements are available. Ideally, we would like the latent transition kernels to be given by the marginal transitions of the Wright-Fisher diffusion from last section,

$$Z_{t_m} | Z_{t_{m-1}}, S \sim \text{WF}(S, N_\text{e}, t_m - t_{m-1}, Z_{t_{m-1}}), \tag{2}$$

where $N_\text{e}$ is estimated as a pre-processing step (Section 6.5). In practise we resort to approximating the distribution in Equation (2) via a Euler-Maruyama scheme (Section 6.4).

Finally, for each $t \in \{t_1, t_2, \ldots, t_T\}$, let $Y_t = (Y_t^1, \ldots Y_t^K)$ denote a noisy observation of the population prevalences at process time $t$. In the single-cell context, this is obtained by counting, for each clone defined in Section 5.1, the number of cells coming from each passage, and normalizing by the number of cells sequenced in that passage. In the bulk sequencing context, see Section 5.1. For simplicity, in both cases we use a normal observation model, i.e., $Y_t^i \mid Z_t^i \sim \mathcal{N}(Z_t^i, \sigma^2_\text{obs})$, where $\sigma^2_\text{obs}$ is an input parameter (values shown in **Supplementary Table 4**).

**6.4 Posterior inference under the fitClone model** Since the marginal distributions of the Wright-Fisher diffusion do not admit closed form expressions, and previous work on exact simulation does not scale to high values of $K$, we resort to discretization using a Euler-Maruyama scheme [23]. For simplicity, we used the same number of grids between all observed time steps. We used a particle Markov chain Monte Carlo (pMCMC) method called Particle Gibbs with Ancestor Sampling and particle rejuvenation [24, 25] to sample from $Z_{t_m}$ and the intermediate Euler-Maruyama steps, and a Metropolis within Gibbs sampler to sample the selection parameters $S$.

**6.5 Estimating the effective population size** Following [26] we use $F_s^{'}$ an unbiased moment-based estimator of the $N_\text{e}$ where $N_\text{e} = \frac{1}{F_S^{'}}$; and $t$ is the number of generations between each passage.

$$F_s^{'} = (1/t)\frac{F_s(1-1/(2\tilde{n})) - 1/\tilde{n}}{(1+F_S/4)(1-1/n_y)} \tag{3}$$

where $F_s = \frac{(x-y)^2}{z(1-z)}$ and $z = (x+y)/2$ and $\tilde{n} = \frac{2n_y n_x}{n_y+n_x}$, the harmonic mean of the sample size (initial population size at the passage) $n_x$ and $n_y$ at the two timepoints. $x$ and $y$ are the minor allele frequencies at the two timepoints.

In the multi-allelic case, we have:

$$F_s = \frac{1}{K}\sum_{i=1}^{K}\frac{(x_i - y_i)^2}{z_i(1-z_i)}$$

This is equivalent to plan 2 in [26], sampling before reproduction and without replacement.

We used the sum of clone sizes as the approximate initial population size at each timepoint/passage. **Supplementary Table 5** lists the resulting $N_\mathrm{e}$ estimates. Since `fitClone` is robust to the choice of $N_\mathrm{e}$ in this range (**Supplementary Fig. 1**C), for set $N_\mathrm{e} = 500.0$ for all datasets analysed in this paper. We note that in our model we assume that the effective population size remains constant over all timepoints. This does not take in account the potential changing population growth rate or the bottleneck effect due to passaging. These phenomena may scale the diffusion time and bias our estimates of evolutionary events including fixation or extinction times. This stretching and compressing of time could be accounted for by adding random effects to the number of generations in the model, for example by taking $\mathrm{N}_{\mathrm{e},t}$ as a piece-wise constant random variable that can vary between passages. This is subject to our ongoing research.

**Supplementary Table 4** shows the other parameters used in the inference over the real datasets.

**6.6 Summarising the posterior distribution** In all cases 10,000 particles and a burn-in equal to 10% of the MCMC samples were used. For the trajectories, we reported $\hat{z}_{1:T}$ where $\hat{z}_t = (\hat{z}_t^1, \ldots, \hat{z}^K)$ and $\hat{z}_t^k$ encodes the post burn-in marginal clonal fraction of clone $k$ at time $t$. As the clonal fraction of the reference clone is not maintained during inference, first for each timepoint $t$, we compute $\hat{z}_t^k = \Sigma_{m=1}^M z_t^k$ for $k \in \{2, \ldots, K\}$ and then we set $\hat{z}_t^1 = 1 - \Sigma_{k=2}^K \hat{z}_t^k$, **Supplementary Fig. 7**A.

The posterior of the selection coefficient vector was summarised by $\hat{s} = (\hat{s}_2, \ldots, \hat{s}_K)$ where $\hat{s}_k$ denotes the post burn-in marginal distribution of clone k, see **Supplementary Fig. 7**B. To compare the selective coefficients of two clones we used a posterior ordering matrix $P_{(K-1)\times(K-1)}$ (**Supplementary Fig. 1**B,C). $P_{i,j} = P(s_i \leq s_j)$ shows the posterior probability that clone i has higher selective coefficient than clone j, with the stronger purple hues (close to 1.0) representing a higher confidence that clone i dominates clone j, and conversely the stronger grey hues (close to 0.0) denote that clone j dominates clone i. Colours closer to white (0.5) represent no dominance. Note that for the lower diagonal elements $P(s_j \leq s_i) = 1 - P(s_i \leq s_j)$ and are omitted for clarity. The diagonal entries are to guide the eyes only.

**6.7 Selecting the reference clone** In our formulation of the Wright-Fisher diffusion one reference clone with selection coefficient of zero has to be chosen. The selective coefficient of the other clones are reported relative to this value. For instance, if the fittest clone is chosen as reference, the other clones will have negative selective coefficients. We chose to set the reference to a clone with an approximately monotonically decreasing trajectory (clonal abundance over time). This choice was motivated by a desire to infer a non-negative value for the fittest clones. **Supplementary Fig. 1**B shows that the model is robust to the choice of the reference clone. We run the inference procedure over the same dataset multiple times, each time changing the reference. The posterior ordering of clones over different choices of clones remained mostly identical.

**6.8 Pairwise relative selective advantage** Let $s_{1:M} = (s_1, s_2, \ldots, s_M)$ be the $M$ post burn-in MCMC samples for the selective coefficients where $s_m = \{s_{m,1}, s_{m,2}, \ldots, s_{m,K-1}\}$ are the sampled selective coefficients of clones 1 to $K-1$ at iteration $m$. Define $P_{i,j} = \Sigma_{m=1}^{M} \mathbb{I}(s_{m,i} > s_{m,j})$ for $i, j \in \{1, \ldots, K-1\}$ be the posterior probability of clone $i$ having a larger coefficient than clone $j$.

**6.9 The deterministic logistic growth model** We developed a closely related population genetics model which incorporates selection via deterministic differential equations, but has closed form solutions. In this model [27], the solution of a deterministic DE, the frequency of each population $c$ out of possible $K$ populations, at time $t$, with selection coefficient $s_c$ is proportional to its starting prevalence $p_0(c)$ multiplied by a power of the relative fitness coefficient $w_c = (s_c + 1)$,

$$f(c, t, s) = \frac{(s_c + 1)^t p_0(c)}{\sum_{k=1}^{K} (s_k + 1)^t p_0(k)} \tag{4}$$

To estimate fitness coefficients $w_{1:T}$ from observed clonal fractions $Y_{1:T}$, we solve the optimization problem in Equation (5) using a limited memory Broyden-Fletcher-Goldfarb-Shanno (BFGS) optimization procedure with box constraints [28]. Note that $f_{t,c}(w)$ is the clonal fraction estimates from Equation (4) and $Y_{t,c}$ is the observed clonal fraction for clone $c$ at time $t$.

$$\min_{w \in \mathcal{W}} (\Sigma_{t=1}^{T} d_t)^{\frac{1}{2}} \tag{5}$$

where $d_{1:T}$ is a vector whose elements are $d_t = (\Sigma_{c=1}^{K} (f_{t,c}(w) - Y_{t,c})^{\frac{1}{2}}$ and $\mathcal{W} = (\mathcal{R} \cup \{0\})^K$ and $w_1 = 1.0$, that is we assume without loss of generality that the index of the reference clone with fitness coefficient 1 is set to 1.

**6.10 Simulation benchmarking** We forward simulated $L = 40$ datasets from the joint distribution of the Wright-Fisher model, with K = 4 (5 clones) and $L = 40$ datasets with K = 10 (11 clones). For each simulated dataset l, we sampled initial clonal abundance vector $Z_{1,l} \sim \text{Dirichlet}(\alpha_{1:K})$ where $\alpha_i = 1$ and selective coefficients from a normal distribution with $s_{i,l} \sim \text{Normal}(0.0, 0.3)$ truncated at $(-0.5, 1)$ for $i \in \{2, \ldots, K\}$ assuming the index of the reference clone is $i = 1$ and $s_1 = 0$. Discretisation constant (step size) $\Delta\tau$ = 0.001 and the standard deviation of the emission model was set to $\sigma_{obs,\text{simul}}$ = 0.001. At simulation and inference, we set $N_{\text{e}} = 500$. The simulation was continued to diffusion time of 0.1 after which 10 equi-distanced samples were recorded as observed values for the process. In all models except the Logistic growth, we put a uniform prior on each component of the s vector, that is, $s_i \sim \text{Uniform}(-5.0, 5.0)$ and a Dirichlet prior on the initial clonal distributions. We set step size $\Delta\tau$ = 0.001, $\sigma_{obs,\text{infer}}$ = 0.01 and use 10,000 particles for 10,000 MCMC iterations. We run 5 different models on the simulated dataset as follows (**Supplementary Fig. 1**): (i) Standard that is the Wright-Fisher model with diffusion approximation. (ii) Logistic growth is the deterministic differential equation Wright-Fisher model. (iii) One-step is the identical to the standard model but applies no discretisation for the time between observations. (iv) Single Clone is the standard model with $K = 1$. See section 6.9 for how logistic model was fit. For the other models, we used the mean absolute error (MAE) of the the post burn-in mean posterior of marginal selection coefficients for each method and the selection coefficient used to generate the simulated data. For each dataset l, the Single Clone model is run K times, once for each clone k, where its input consists in the observed clonal fraction of only one clone. For this model we report the averaged MAE across clones per dataset. The single clone model performs worst which suggests that treating each clone independently is sub-optimal.

**6.11 Software and implementation** The software implementation of `fitClone` is available at: [`https://github.com/UBC-Stat-ML/fitclone`]

### 7 scRNAseq data analysis

**7.1 Quality control** Count matrices were generated using CellRanger version 3.1.0. Cells were considered to have passed a quality control filter (QC-filter) and retained for subsequent analysis if they met the following criteria: (i) at least 1000 genes detected, (ii) less than 20% of counts (UMIs) mapping to genes from the mitochondrial genome ("mitochondrial genes"), (iii) fewer than 60% of counts (UMIs) mapping to ribosomal genes, and (iv) the total counts (UMIs) per cell was at most 3 median absolute deviations lower than the overall median. Cells not matching all criteria were filtered using the `calculateQCMetrics` and `isOutlier` functions in the `scater` package [29]. Then, all the mouse cells were eliminated. A cell is called a mouse cell if the total number of counts in a mouse alignment of the 10x sample was greater than the total number of counts in a human alignment. Finally, we eliminated doublets using package `scrublet`[30] (**Supplementary Table 2**).

**7.2 Gene expression normalization** Sample level normalized log expression values were computed using `scran` [31] with grouping variables calculated from clustering using the `quickCluster` function [32]. Overall normalized expression was computed by merging sample level matrices and recalculating normalized expression with grouping labels computed on the merged matrix. Batch correction between libraries was performed using Scanorama [33] on the overall normalized expression values with default parameters and the library id as the batch label.

**7.3 Dimensionality reduction** Principal component analysis was performed on the batch corrected expression matrix using the `scater` [29] R package. A single two dimension t-SNE embedding was generated using the first 50 principle components. This embedding was used in downstream analysis and visualizations.

**7.4 Pathway Enrichment Networks** Enriched pathways were computed from differentially expressed genes (adjusted p-value $< 0.01$) ranked by log fold change. A normalized enrichment score (NES) was calculated from a ranked gene set enrichment analysis (GSEA) [34] performed on each subset of differentially expressed genes using the hallmark gene set collection from MSigDB [35]. Significantly enriched pathways (adjusted p-value $< 0.01$) and pathway specific differentially expressed genes were included in network enrichment figures. Pathway nodes were colored by NES value. Edges are defined between pathways sharing genes. All analysis and visualization was performed using `gseapy` and `networkx` [36] Python packages.

**7.5 Integrative genome-transcriptome analysis with `clonealign`** `clonealign` version 1.99.2 was used to align scRNAseq cells to the DLP+ clones obtained. First, genes whose purity was less than 60% in any clone (40% for *X5 UT*) were removed, where the purity of a gene in a clone is defined as the percentage of cells that have the modal copy number for that gene and clone. Then, we used the following clonealign parameters: `n_repeats = 3`, `mc_samples = 1`, `learning_rate = 0.07`, `max_iter = 500`, `data_init_mu = TRUE` for all samples except `FALSE` for *X5 UTU*, `data_init_mu = FALSE`, `saturation_threshold = 6` and `clone_call_probability = 0.9`.

**7.6 Differential expression analysis** Differential expression quantifiers including log2 fold change and FDR were computed using the R 3.6.0 Bioconductor package `edgeR_3.26.0` that implements scRNAseq differential expression analysis methodology based on the negative binomial distribution. Given the raw counts for the cells in two clones, we first call the `estimateDisp()` function to estimate the dispersion by fitting a generalized linear model that accounts for all systematic sources of variation. Next, we use the `edgeR` functions `glmQLFit()` and `glmQLFTest()` to perform a quasi-likelihood dispersion estimation and hypothesis testing that assigns false discovery rate values to each gene. In the track and volcano plots, a positive log2 fold change value for clone X relative to clone Y signifies that the gene is significantly more upregulated (at a given FDR threshold) in X than in Y while taking into consideration all the expression values for all the genes in both clones. Similarly, a gene with negative log2 fold change is significantly more downregulated in X than in Y.

**7.7 Phenotypic volume analysis** Phenotypic volume [37] was computed for each of the 13 scRNAseq TNBC PDX libraries. After removing the remaining mitochondrial and ribosomal genes, we selected 1,983 common genes that were detected in at least 200 cells in each of the 13 libraries. Then, we sampled uniformly at random 800 cells from each library, thus yielding a 1,983 genes x 800 cells matrix of normalized log2 counts for each library. Next, we computed the covariance matrix such that only the common values between every pair of genes was considered, while the missing values were ignored. The phenotypic volume is the sum of log10 of all the singular values of the covariance matrix. We repeated the entire process 20 times.

**7.8 RNA velocity analysis** We used `velocyto.R_0.6` [38] to estimate RNA velocities. Briefly `run10x` from velocyto CLI was used to preprocess the cellranger output directories from the 10x Genomics platform for each timepoint of the TNBC Rx arm individually. The resulting loom files were merged using the `combine` command in the `loompy` package as recommended by the authors. Cells and genes were selected as in section 7.7. RNA velocities were computed using the `gene.relative.velocity.estimates(emat, nmat, deltaT=1, fit.quantile = 0.02, kCells = 20)` routine with an identical embedding matrix as in the main text. Absolute cumulative RNA velocity was defined as the mean average magnitude of the estimated velocities of all genes per cell.

### 8 Supplementary figures

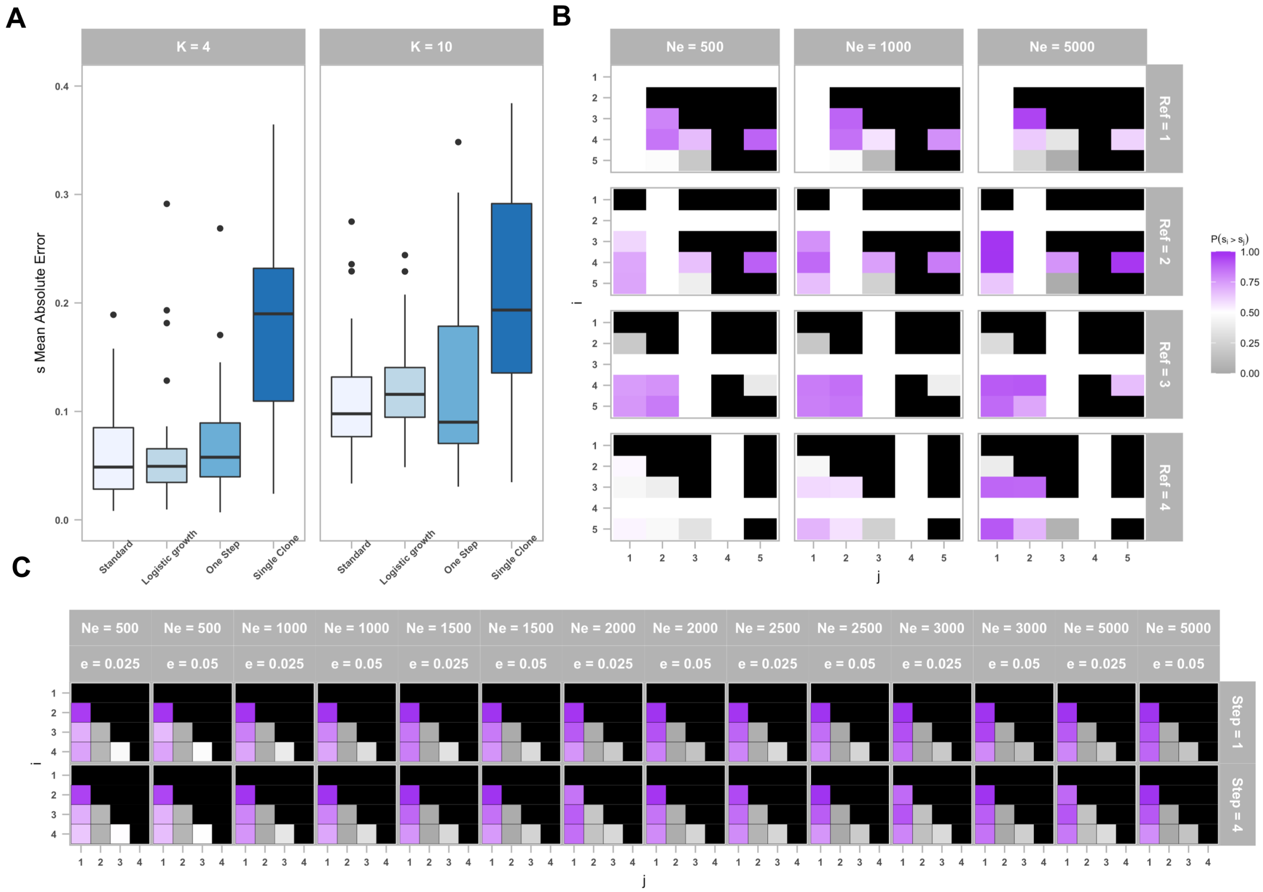


**Supplementary Figure 1** Simulation studies for the `fitClone` model. **A)** Comparison to baseline methods for K = 4 clones (left) and K = 10 clone (right) **B)** Posterior ordering of clones based on their inferred posterior selective coefficients across three values of effective population size (columns) and different choice of the reference clone (rows) **C)** Posterior ordering of clones across different hyper parameters in the `fitClone` model. Effective population size in the range of 500 to 5,000 (top column), and observation error (bottom column). Minimum number of interpolations between two observation (rows)

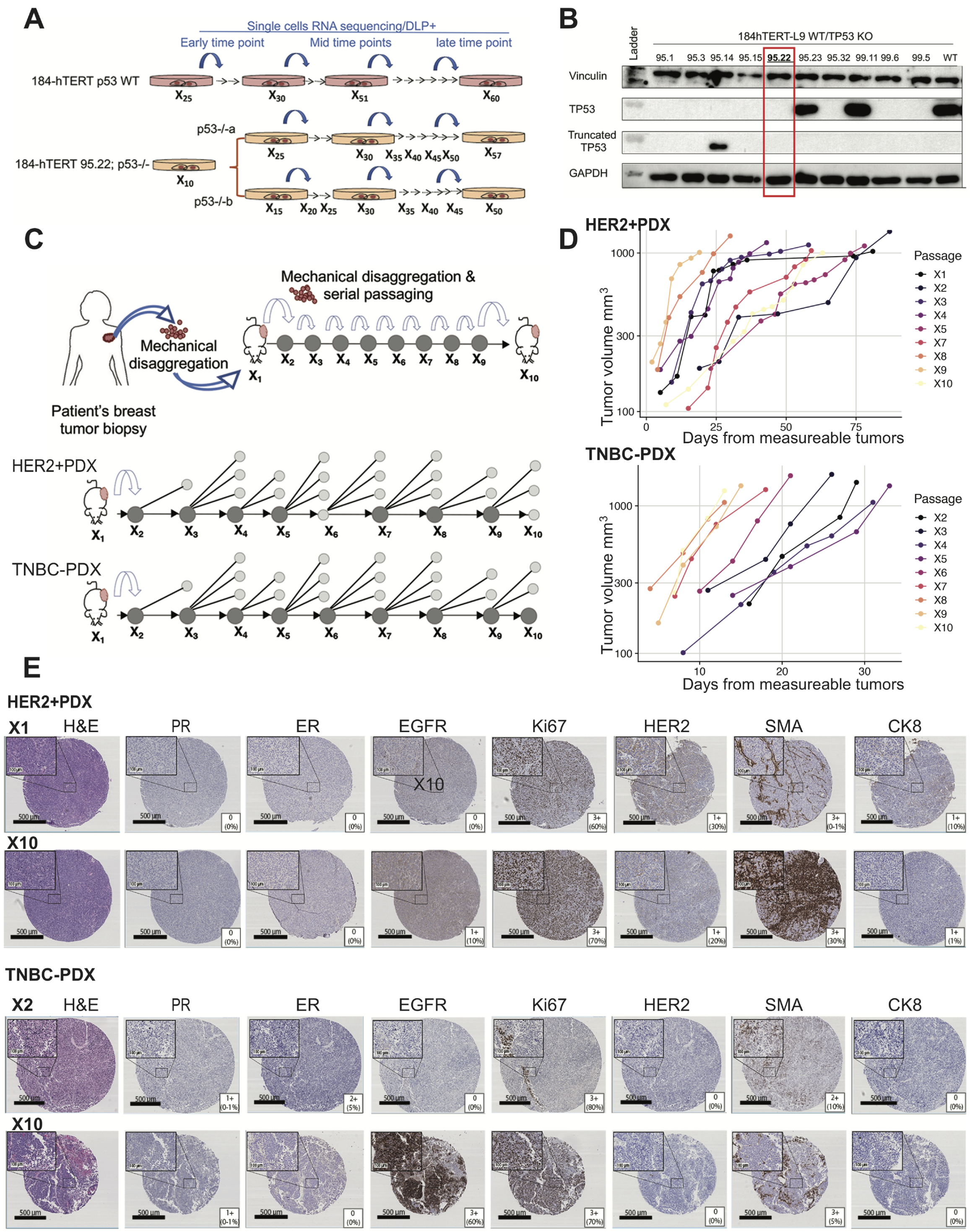


**Supplementary Figure 2** Overview of experimental design and PDX growth curves. **A)** Top: Serially passaged 184-hTERT L9 WT cell line; Bottom: 184-hTERT L9 95.22; p53 null cell line by CRISPR technology. Parallel branches P53$^{-/-}$ a and P53$^{-/-}$ b were derived from the same tenth passage. **B)** Western blot confirming knock out of TP53 from 184-hTERT WT cell line. Clones shown along top. **C)** Top: Schematic for PDX timeseries; Bottom: Serial sampling of HER2+ and TNBC PDX tumours; Dark grey circles represent each sampled mouse for scWGS. The light grey circles representing the replicates of tumour-bearing mice at the same timepoint. **D)** Individual tumour growth from each passage of TNBC and HER2+ PDXs. **E)** IHC of HER2+ and TNBC tumours at early and late passages, 4x and 20x (insets). Scale bars 500 µm and 100 µm (insets). Antibodies and TMA scores (**Supplementary Tables 6** and **7**).

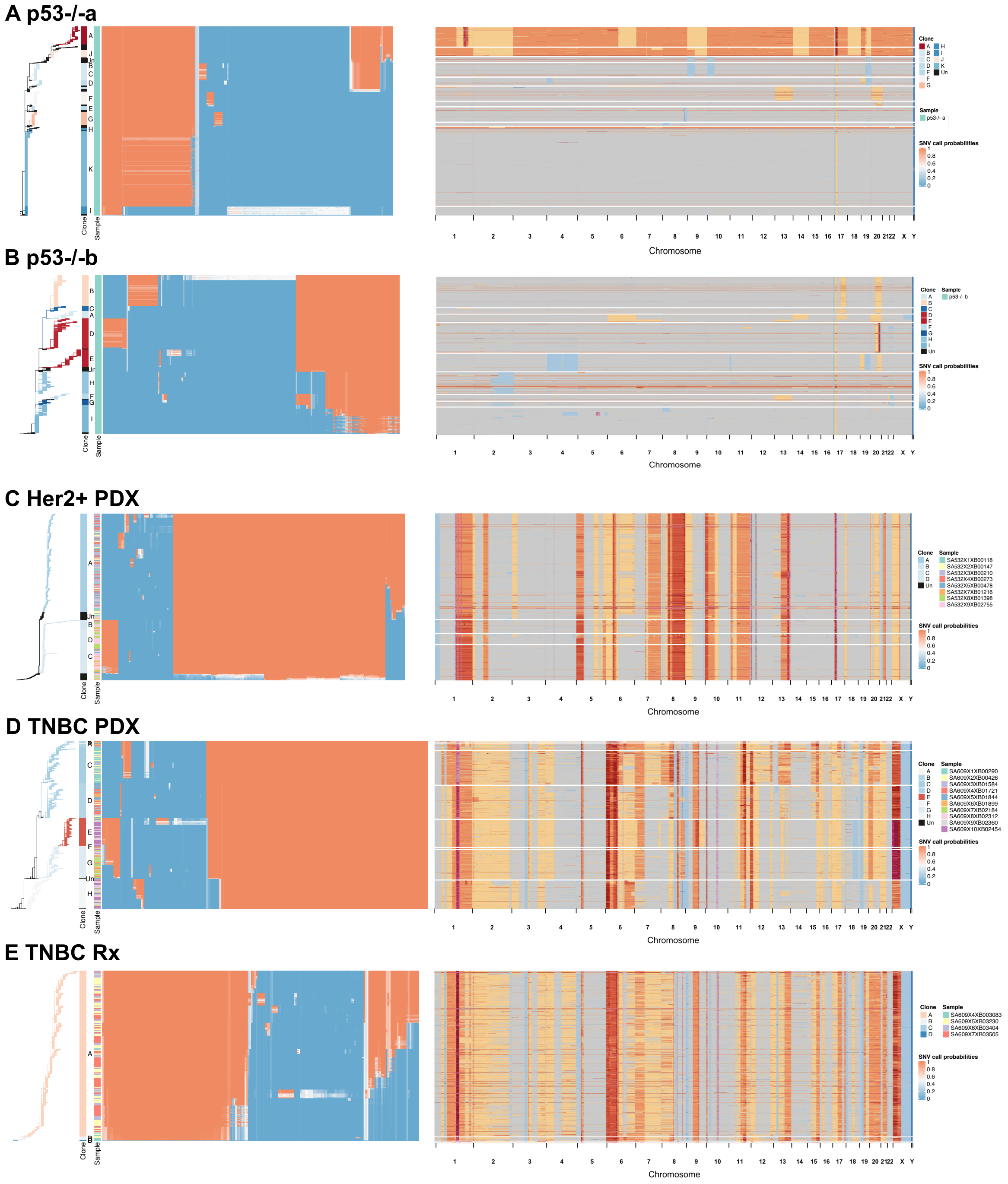


**Supplementary Figure 3** Single nucleotide variant (SNV) and Copy number profiles of the cell lines and PDX libraries. **A)** Phylogenetic tree with clone labels next to SNV and copy number profile heatmaps for *p53-/-a*. Rows are individual cells and columns are SNVs (left) and copy number alterations (right). For SNVs, orange indicates high probability while blue indicates low probability. For copy number, legend encoding copy number colours is shown (right). **B-E)** Similar to **A**) but for *p53-/-b*, HER2+, TNBC, and TNBC Rx respectively

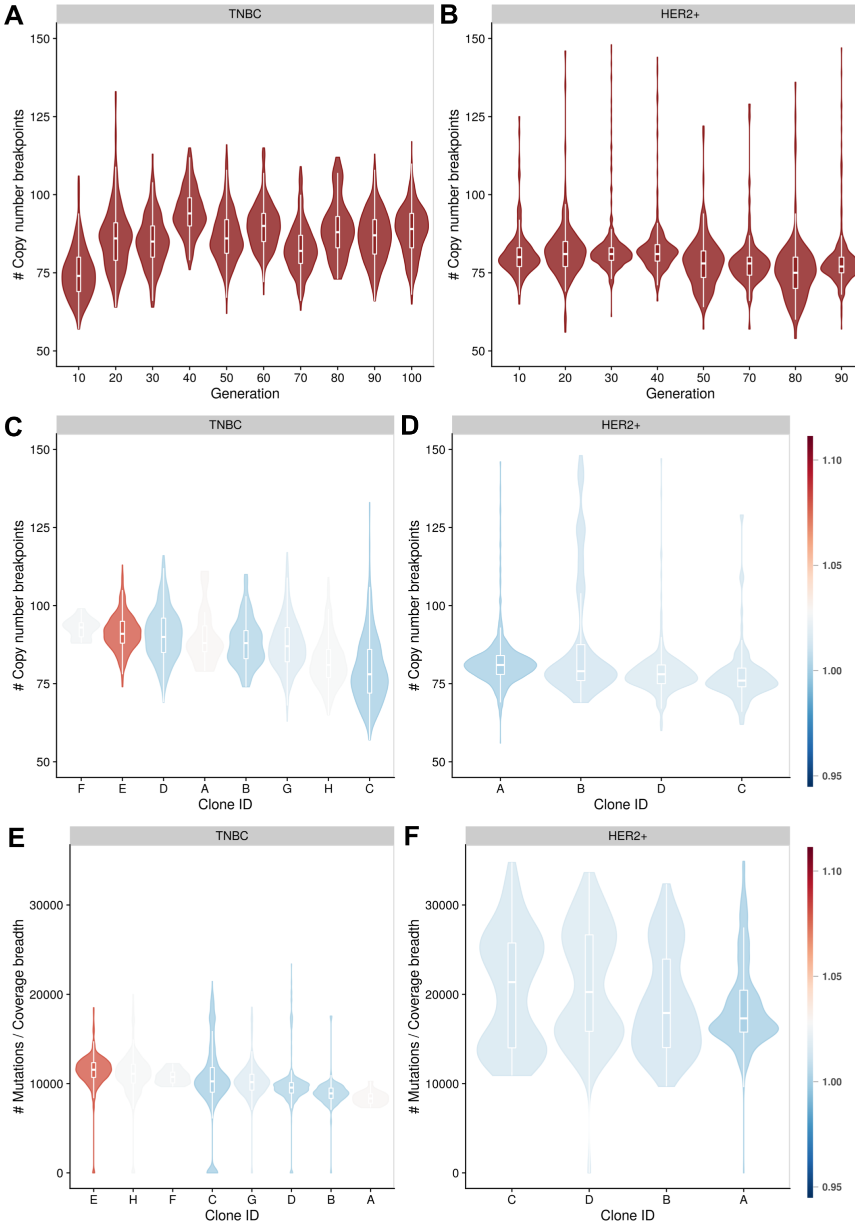


**Supplementary Figure 4** Structural variant and mutation rates of PDX lines. Distribution over copy number breakpoints/cell as a function of generation for **A)** TNBC **B)** HER2+. Clone specific distributions over copy number breakpoints/cell, colored by fitness coefficients for C) TNBC **D)** HER2+. Clone specific distributions over point mutations/cell, colored by fitness coefficients for **E)** HER2+ and F) TNBC

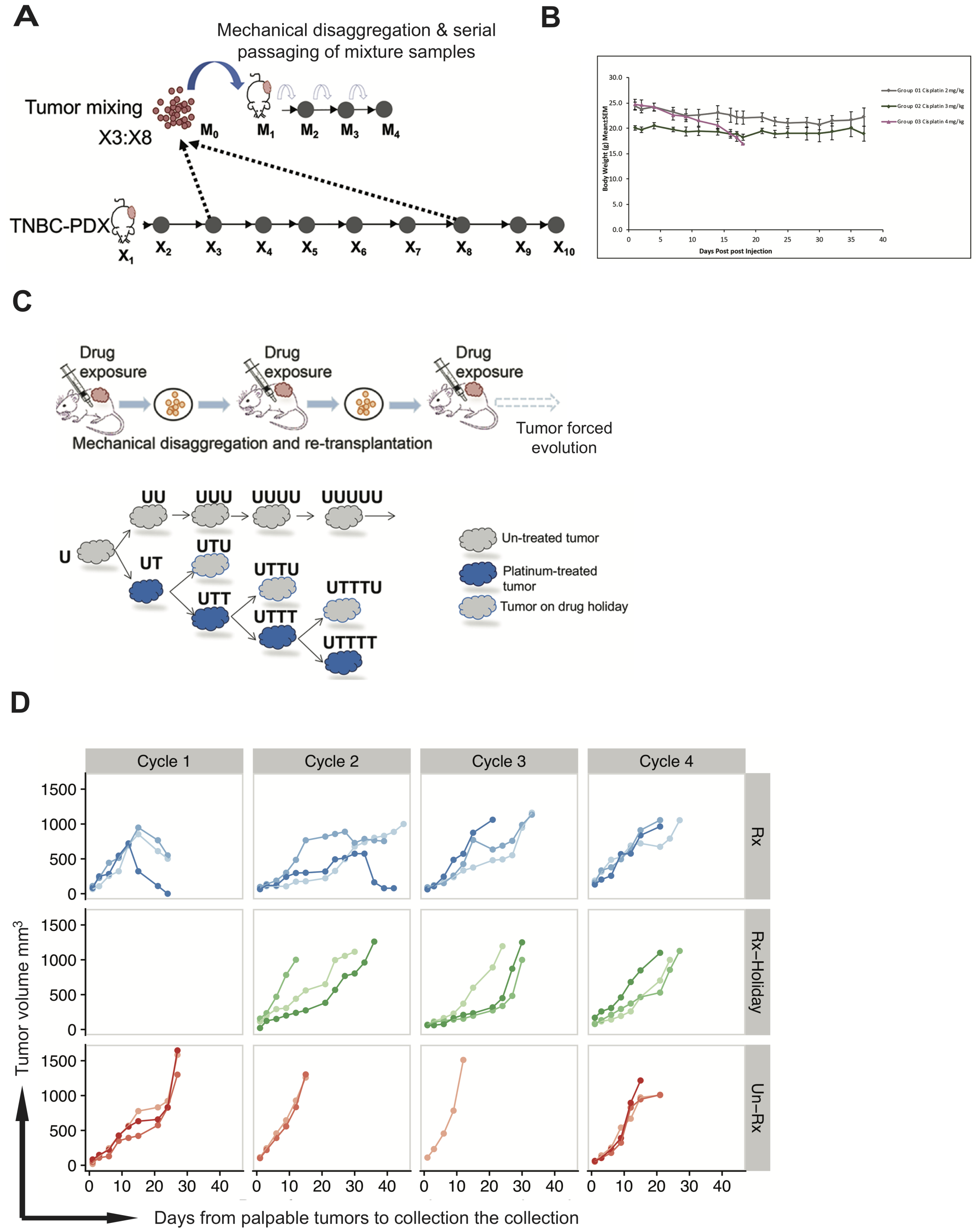


**Supplementary Figure 5** Fitness validation with tumour mixing and drug perturbation. **A)** Schematic overview of clonal mixture experiment showing source samples from the original timeseries and serial propagation into a new line. **B)** Mouse body weight graph recorded during maximum tolerated dose (MTD) evaluation of cisplatin in NRG mice (n=3 in each study cohort). **C)** Experimental design of cisplatin treatment in PDX.The residual tumour from one treated mouse was re-transplanted in the next (n=4). The solid blue colour representing cisplatin treated tumours *(UT,UTT,UTTT,UTTTT)*; blue outlined in grey represent drug holiday *(UTU,UTTU,UTTTU)*. Grey represent the untreated series *(U,UU,UUU,UUUU,UUUUU)*. **D)** Tumour response curves in each cycle of cisplatin treatment.

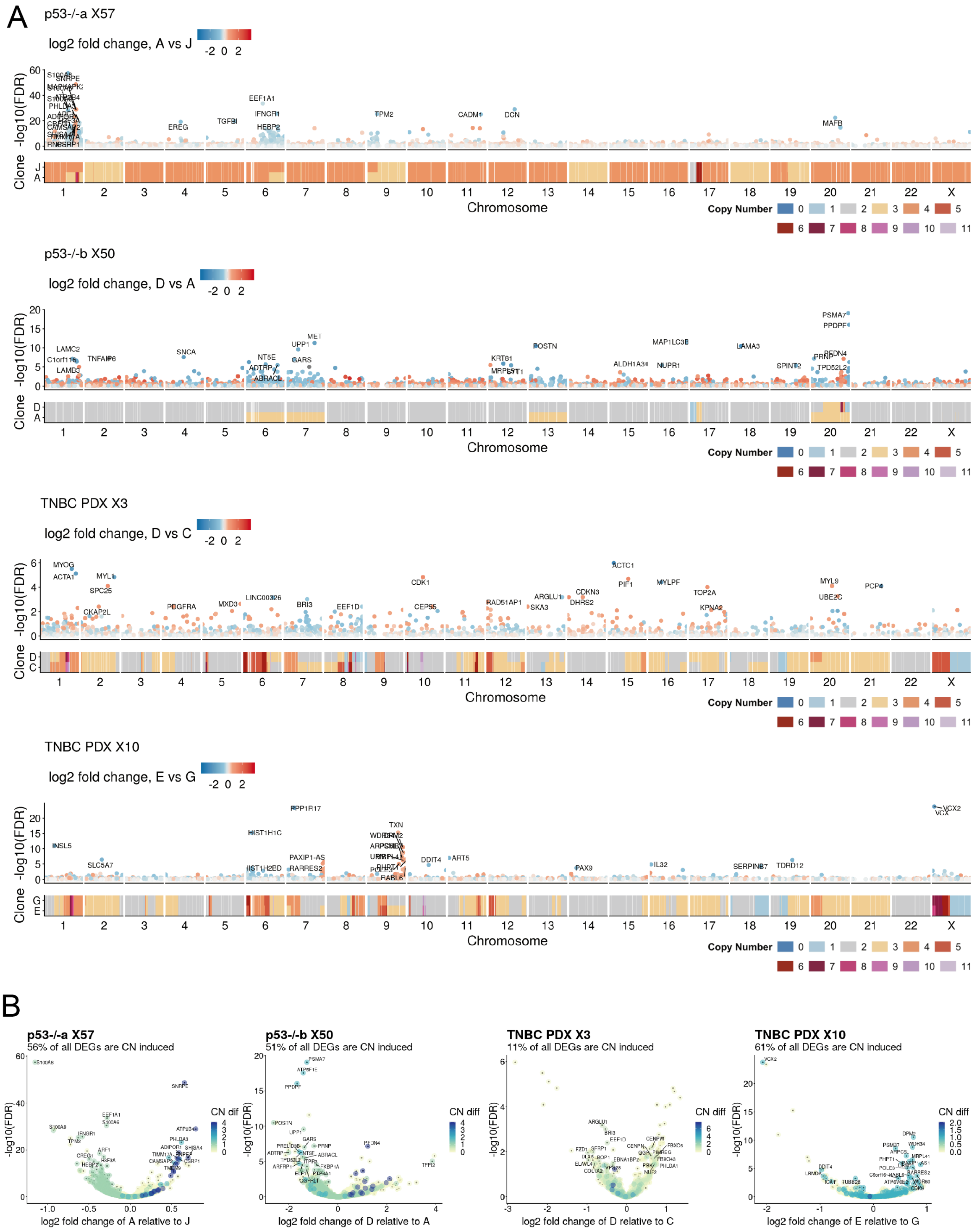


**Supplementary Figure 6** Pairwise clone-specific differential gene expression genome-wide. **A)** Track plots for *p53-/-a X57* clone A versus J (tetraploid clones); *p53-/-b X50* clone D versus A; TNBC *PDX X3* clone D versus C; and TNBC *PDX X10* clone E versus G. **B)** Volcano plots corresponding to the track plots in panel **A**

.

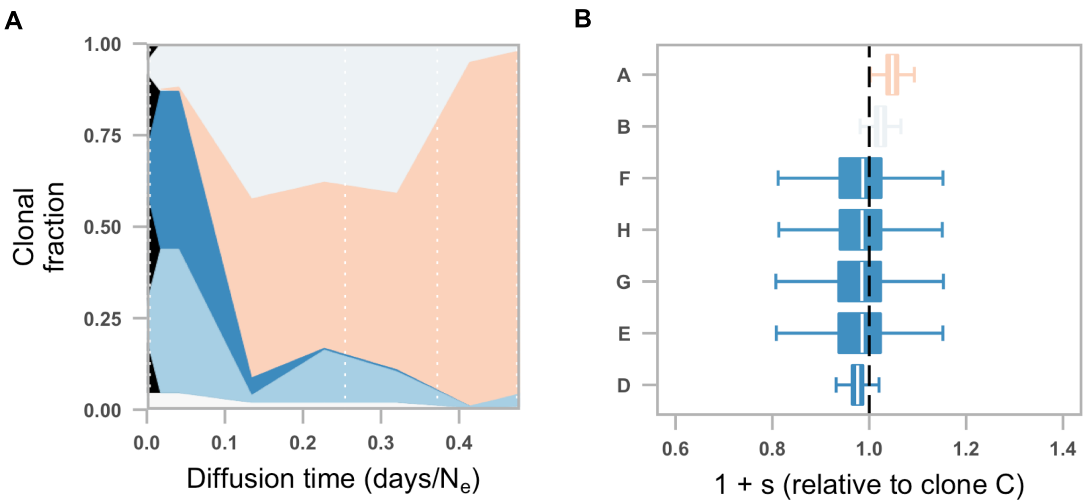


**Supplementary Figure 7** Fitness analysis of the TNBC Rx line. **A)** The inferred trajectories **B)** Marginal posterior distribution of selection coefficients of each clone, sorted by their mean selective coefficients

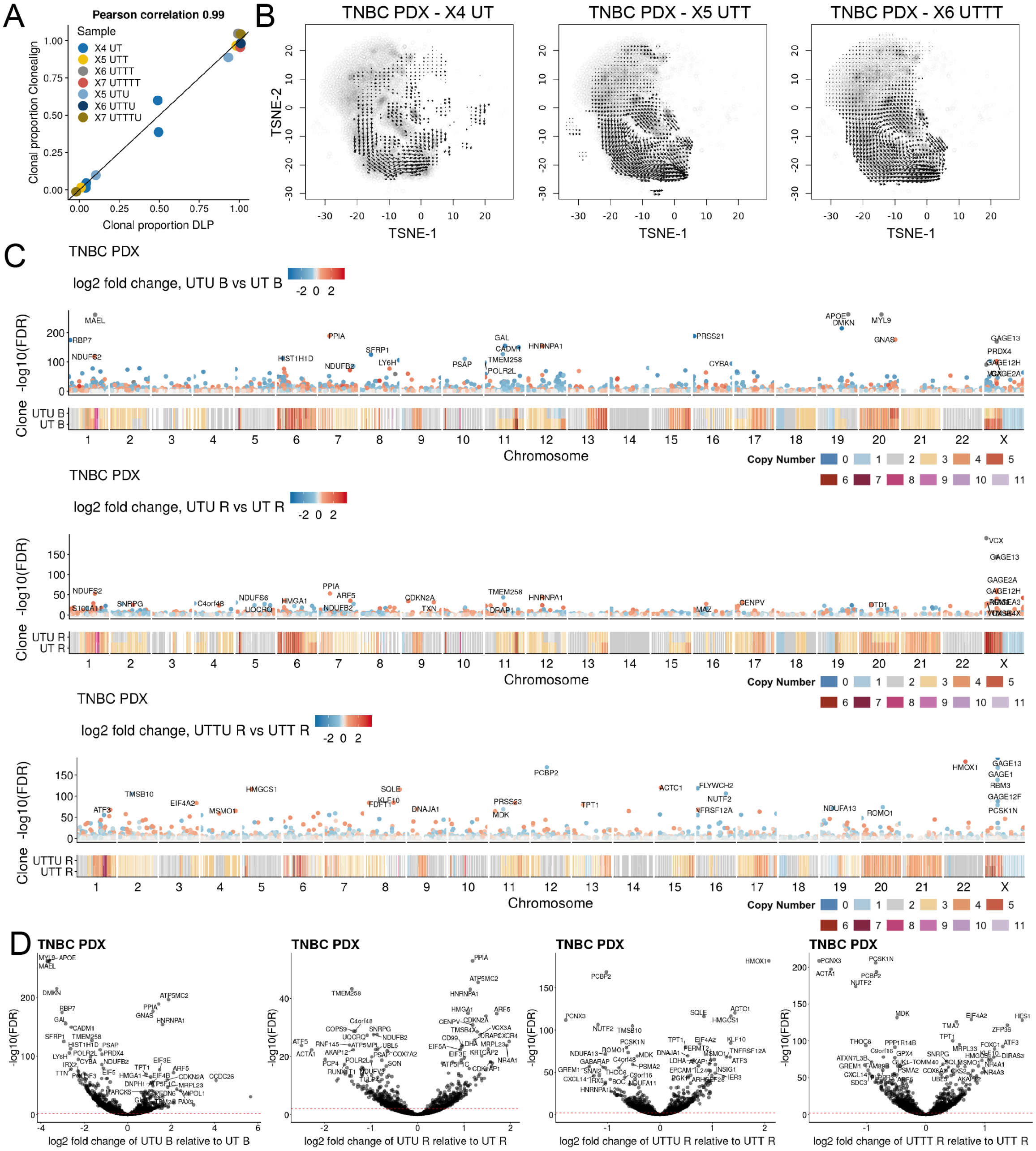


**Supplementary Figure 8** Phenotypic diversity following cisplatin treatment in *TNBC PDX* treated and holiday lines. **A)** Plot showing close to 1 Pearson correlation between the DLP+ clone proportions and the clonealign clones. **B)** Visualization of transcriptional velocity vectors for the X4-X6 Rx passages. **C)** Track plots showing phenotypic diversity of the same clone (B or R) in holiday sample versus the previous treated timepoint. **D)** Volcano plots corresponding to the track plots in panel **B)**, and an additional volcano plot (right) comparing clone R at X6 vs X5 treated timepoints

.

### 9 Supplementary tables

**Supplementary Table 1** DLP+ libraries generated for p53 WT, p53-/-a, p53-/-b, HER2+ and TNBC timeseries indicating timepoint, number of cells (initially and after different QC levels), median reads per cell and quality metrics.

**Supplementary Table 2** 10x scRNAseq libraries indicating timepoint, initial number of cells, number of cells after three quality control measures and mean, standard deviation and median of counts in the entire library (entries with counts $< 100$ were eliminated from the mean, standard deviation and median metrics).

**Supplementary Table 3** Clones and their respective 1+s coefficients inferred for WGS-bulk-TNBC , p53 WT, p53-/-a, p53-/-b, HER2+ and TNBC timeseries indicating timepoint, clonal abundance (frac), number of cells (if applicable), and mean, median and standard deviation of selective coefficients.

**Supplementary Table 4** Real data parameters

| | Dataset | $\epsilon$ | runtime | niter | $\Delta\tau$ | $\sigma_p$ | avg_ESS | min_ESS | rejection_rate |
|---|---|---|---|---|---|---|---|---|---|
| 1 | *p53 WT* | 0.01 | 2:57:22. | 10,000 | 0.05 | 0.05 | 911.90 | 448.19 | 0.77 |
| 2 | *p53-/-a* | 0.01 | 35:18:46. | 100,000 | 0.05 | 0.03 | 3209.54 | 506.66 | 0.61 |
| 3 | *p53-/-b* | 0.03 | 31:29:55. | 100,000 | 0.04 | 0.05 | 2077.27 | 906.95 | 0.89 |
| 4 | HER2+ | 0.01 | 48:57:27. | 100,000 | 0.05 | 0.03 | 26303.53 | 25299.69 | 0.38 |
| 5 | TNBC | 0.03 | 40:05:32. | 100,000 | 0.05 | 0.05 | 1379.81 | 223.08 | 0.91 |
| 6 | TNBC-mixture | 0.01 | 29:13:01. | 100,000 | 0.04 | 0.05 | 3745.76 | 1268.09 | 0.73 |
| 7 | TNBC-Rx | 0.03 | 30:05:33. | 100,000 | 0.10 | 0.05 | 4362.86 | 2768.55 | 0.73 |
| 8 | WGS-bulk-TNBC | 0.012 | 35:12:29. | 100,000 | 0.10 | 0.025 | 14560.09 | 11754.76 | 0.71 |

**Supplementary Table 5** Parameters per dataset. Effective population size estimates and number of loci used in the SNV analysis

| | Dataset | Ne | Number of SNVs |
|---|---|---|---|
| 1 | *p53 WT* | 985.79 | NA |
| 2 | *p53-/-a* | 614.93 | 6,645 |
| 3 | *p53-/-b* | 422.14 | 7,905 |
| 4 | HER2+ | 1,461.70 | 40,309 |
| 5 | TNBC | 468.30 | 33,178 |
| 6 | TNBC-Mixture | 177.01 | NA |
| 7 | TNBC-Rx | 333.34 | 25,685 |

**Supplementary Table 6** List of antibodies and experimental conditions. Summary of antibodies clones and their suppliers used for staining TMAs for IHC and performing Western blots for p53 WT and $P53^{-/-}$. **(*RTU: Ready to use; N/A: not applicable)**

| | Specimen | Stain/IHC | Vendor / Ab Clone | Dilution |
|---|---|---|---|---|
| 1 | TMA1 & 2 | CK14 | Empire Genomics clone LL002 | 1 in 50 |
| 2 | TMA1 & 2 | Ck5/6 | Dako D5/16 B4 | *RTU |
| 3 | TMA1 & 2 | Ck8 (CAM5.2) | BC Bioscience CAM5.2 | 1 in 10 |
| 4 | TMA1 & 2 | EGFR | Epitomics 1902-1 | 1 in 100 |
| 5 | TMA1 & 2 | ER | Ventana clone SP1 | *RTU |
| 6 | TMA1 & 2 | H&E | *N/A | *N/A |
| 7 | TMA1 & 2 | INPP4B | Abcam EPR3108Y ab81269 | 1 in 50 |
| 8 | TMA1 & 2 | Ki67 | Abcam ab16667 | 1 in 400 |
| 9 | TMA1 & 2 | PR | Abcam ab30285 | 1 in 200 |
| 10 | TMA1 & 2 | Slug/Snail | Abcam ab85936 | 1 in 125 |
| 11 | TMA1 & 2 | SMA | Dako clone 1A4 | 1 in 100 |
| 12 | TMA1 & 2 | Trichrome | Sigma-HT15-1KT | *N/A |
| 13 | TMA1 & 2 | Twist | NB120-49254 | 1 in 200 |
| 14 | TMA1 & 2 | Vimentin | Dako V9 | RTU |
| 15 | TMA1 only | E-Cad | Cell Signal 3195 | 1 in 100 |
| 16 | TMA1 only | HER2 | Roche 4B5 | 1 in 8 |
| 17 | TMA2 only | E-Cad | Dako NCH-38 | *RTU |
| 18 | TMA2 only | HER2 | Ventana clone 4B5 | *RTU |
| 19 | Western Blot | GAPDH | Santa Cruz (1-19) | 1 in 500 |
| 20 | Western Blot | TP53 | Santa Cruz (DO-1) | 1 in 500 |
| 21 | Western Blot | Vinculin | Santa Cruz (H-300) | 1 in 500 |

**Supplementary Table 7** IHC Scores of un-treated and treated a time series tissue microarray (TMA). HER2+ and TNBC PDX serial passages and cisplatin treated serial passages on TMA with scores of various stains

**Supplementary Table 8** Pathway enrichment analysis scores.
